# Capsid structures of Atlas viruses reveal an engineerable nanoparticle scaffold

**DOI:** 10.64898/2026.03.22.711591

**Authors:** Haoming Zhai, Yangci Liu, Oscar Beechey-Newman, Yorgo Modis

## Abstract

Endogenous viruses and retroelements occupy a large fraction of genomes, but the properties of proteins they encode remain largely uncharted. Here, we define Atlas viruses, a new clade of endogenous viruses in nematodes with a retrovirus-like genome organization, phlebovirus-like envelope glycoproteins, and highly divergent Gag proteins. Cryo-EM structures of particles from three Atlas viruses reveal thin-shelled icosahedral assemblies containing 60, 240, 420 or 720 capsid proteins and ranging from 20 to 60 nm in diameter. The capsid fold and assembly interactions are distinct from those of orthoretroviruses but reminiscent of spumaretrovirus and dArc capsids. Atlas capsids package nucleic acids, enter cell by endocytosis, and disassemble at endolysosomal pH. Through structure-guided engineering, we generated recombinant Atlas particles with increased pH resilience and particles displaying a CTLA-4 receptor-targeting tag. This work identifies Atlas capsids as attractive candidates for development as engineerable scaffolds for tissue-specific nucleic acid delivery.

## INTRODUCTION

Retroviruses and retroelements encode reverse transcriptases, which convert their RNA genome into DNA that can integrate it into host chromosomes^1,2^. When integration occurs in the germline, viral sequences become vertically inherited. Such endogenized viral elements make up 15% of the human genome and serve as a genetic reservoir from which new genes and regulatory elements can evolve^3–6^. Virus-derived proteins were co-opted to fulfill vital functions at various points in evolution. In plants and lower eukaryotes, gamete fusion is driven by homologs of class II viral envelope glycoproteins (Envs)^7,8^. In mammals, Envs co-opted from endogenous retroviruses drive trophoblast fusion during placental development^9,10^. In nematodes, the retroelement-derived Cer1 capsid packages pathogen-derived RNA into virus-like particles that transmit pathogen-avoidance behavior across generations^11^. A similar RNA transfer mechanism occurs in mammalian neurons, where Arc, a repurposed retroelement capsid, transports RNA in virus-like particles across synapses, contributing to neuronal plasticity and memory formation^12–14^.

Newly available whole-genome sequences and improved tools for analyzing intergenic sequences have revealed thousands of protein-coding genes derived from endogenous viral elements. Many of these have the hallmarks of repurposing, or domestication: conservation across species, evidence of purifying selection, and tissue-specific expression, particularly in reproductive and nervous tissues^15,16^. Two genes encoding capsid proteins of retroviral origin, PEG10 and RTL1, have been co-opted in mammals^15,17^ and are essential for embryonic development in mice^18,19^. PEG10 and PNMA2, another retroviral capsid protein, form virus-like particles that package mRNAs and deliver them into cells, identifying them as candidate delivery vehicles for RNA-based therapies^20,21^.

Efficient nucleic acid delivery remains a major bottleneck for gene and cell therapies. Lipid nanoparticles are difficult to target to tissues beyond the liver^22^. Adeno-associated virus (AAV) vectors have enabled major advances in gene delivery, but their small cargo capacity, immunogenicity, and persistent transgene expression constrain certain applications^23,24^. Retroviral vectors are difficult to manufacture at scale and prone to off-target genomic insertion^25,26^. These limitations are particularly relevant for CRISPR/Cas9-based gene editing approaches and emerging *in vivo* chimeric antigen receptor T-cell therapies, which require large nucleic acid payloads and cell type-selective targeting^27,28^. Recombinant viral capsid proteins are being explored as nanoparticle scaffolds that can be recombinantly produced, structurally engineered, and functionalized for targeting without the need for viral replication or integration machinery^20,21^. However, the structures, assembly principles and engineering potential of most endogenous viral capsid proteins remain unknown. Charting this structural space could reveal new classes of protein assemblies capable of overcoming limitations of currently available nucleic acid delivery systems.

We previously identified Atlas virus in the hookworm parasite *Ancylostoma ceylanicum* as an endogenous belpaovirus with a unique combination of retroviral and non-retroviral features^29^. Widespread in metazoans, belpaoviruses have retrovirus-like genome organizations and belong to the order *Ortervirales*^30,31^. However, instead of a retroviral Env, Atlas virus encodes proteins homologous to phlebovirus G_N_ and G_C_ glycoproteins^32^, including a functional class II membrane fusion protein not previously described in a reverse-transcribing virus^29^. The Atlas Gag proteins are highly sequence-divergent relative to canonical retroviral Gags. Here, we identify a new clade of belpaoviruses in nematodes with phlebovirus-like glycoproteins and Atlas-like capsid domains. We report cryo-EM structures of capsid particles from Atlas virus and two other clade members. The structures reveal thin-shelled icosahedral assemblies structurally distinct from orthoretrovirus capsids but reminiscent of spumaretrovirus^33^ and *Drosophila* dArc capsids^13^. Atlas-clade capsid particles assemble without the canonical retroviral major homology region, package nucleic acids, are endocytosed, and disassemble at endolysosomal pH.

Moreover, we show that Atlas capsids can be engineered to modulate particle stability or display a heterologous targeting module. Our work identifies Atlas capsid particles as divergent viral assemblies and provides a structural framework for future engineering of novel nucleic-acid delivery scaffolds.

## RESULTS

### A set of nematode Atlas viruses with class II Env and distinctive Gag proteins

Atlas is an intact viral element, flanked by 100% identical 271-nucleotide long terminal repeats (LTRs). It encodes a 2,828-amino acid Gag-Pol-Env polyprotein^29^. Structural modeling of Atlas Gag with AlphaFold 3^34^ yielded three structured, mostly α-helical domains separated by unstructured linkers (**Fig. 1a,b**). Following the nomenclature for most *Ortervirales* we refer to these domains as matrix (MA, residues 2-137), capsid (CA, residues 188-372) and nucleocapsid (NC, residues 412-487). However, Atlas MA and CA have no significant sequence similarity to retroviruses MA and CA. The predicted structure for MA was a four-helix bundle. AlphaFold 3 predicted a dimeric assembly for MA with high confidence (ipTM = 0.82), but not a trimeric assembly (ipTM = 0.35). The predicted antiparallel dimer did not bear any similarity to reported structures of retrovirus matrix dimers^35–37^. Moreover, Atlas MA lacks the N-myristylation signal responsible for membrane targeting of most orthoretroviruses.

**Figure 1.**
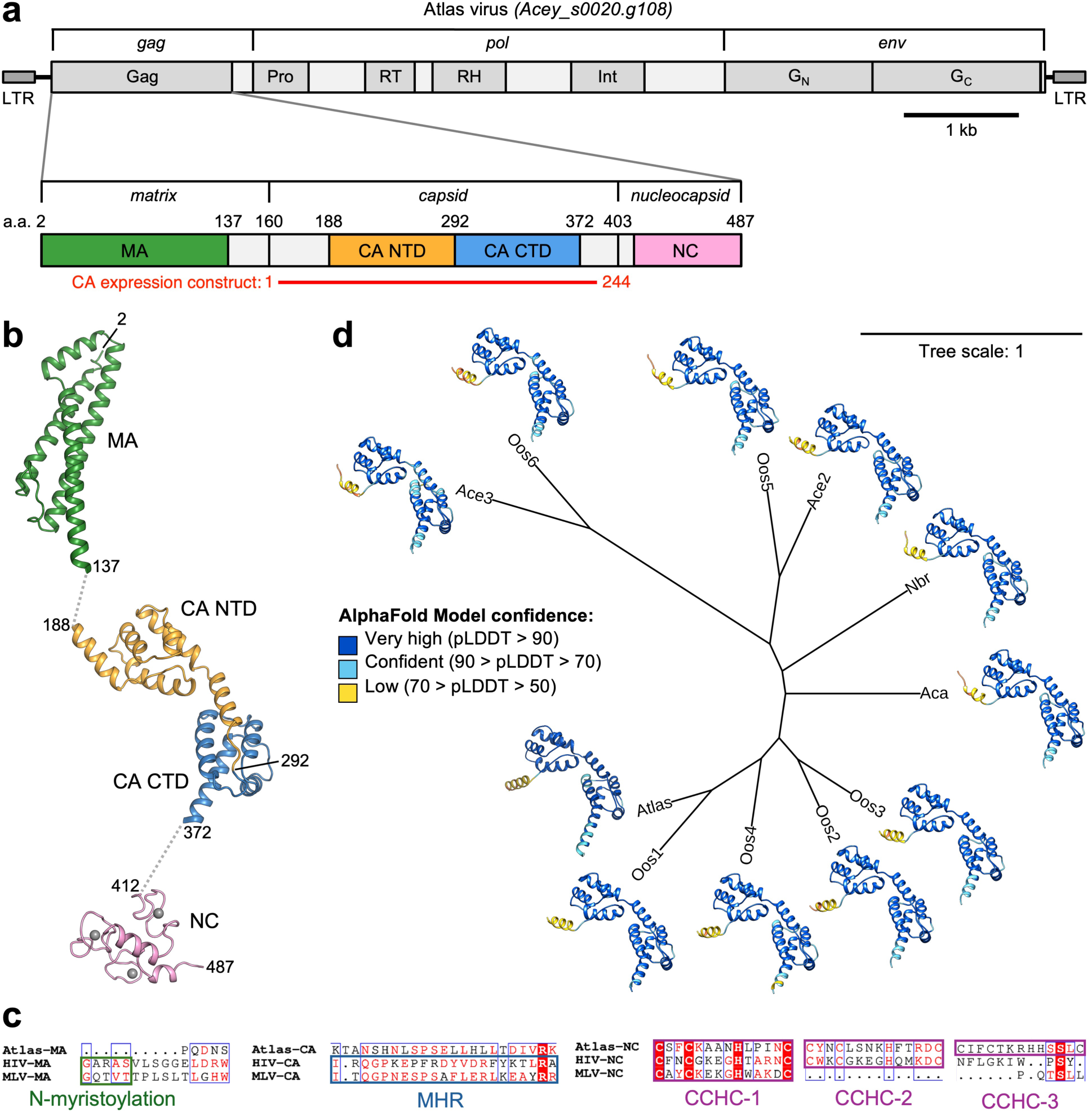
Genetic architecture and phylogenetic definition of the Atlas virus clade. **(a)** Genetic architecture of Atlas virus (GenBank: EPB78661). MA, matrix. CA, capsid. NC nucleocapsid. NTD, N-terminal domain. CTD, C-terminal domain. **(b)** AlphaFold 3 models of Atlas MA, CA and NC. Disordered regions (predicted Local Distance Difference Test (pLDDT) confidence score < 70) are not shown. **(c)** Extract from a multiple sequence alignment of Atlas virus and two model retroviruses, HIV-1 and murine leukemia virus (MLV), showing the retroviral N-myristoylation sites at the Gag N-terminus, major homology region (MHR) in CA, and cystine/histidine zinc knuckles in NC. There is no significant sequence identity between the Atlas and retrovirus MA and CA domains. **(d)** Phylogenetic tree and AlphaFold 3 models of the set of 11 viruses identified here as the Atlas virus clade. Generated with iTOL v7^51^.

Atlas CA was predicted to consist of two independently folded domains, CA_NTD_ (residues 188-291) and CA_CTD_ (residues 292-372) (**Fig. 1b**). Notably, Atlas CA lacks the major homology region (MHR) conserved in most retroviruses (**Fig. 1c**)^38^. Atlas NC is predicted to contain three CCHC zinc knuckle motifs, a distinguishing feature of belpaoviruses, whereas retrovirus NCs typically contain one or two zinc knuckles^39,40^.

To determine whether any other viruses shared the distinctive genetic and structural features of Atlas virus, we implemented a two-step BLAST pipeline. First, the full-length Atlas polyprotein was used as the query in a PSI-BLAST search against the non-redundant protein database. Only hits with >70% alignment coverage were retained (25 hits) to select for sequences encoding a phlebovirus G_N_-G_C_-like Env. Second, this filtered set was rescreened with BLASTp using as the query Atlas CA a.a.188-372, referred to hereafter as the CA core. We retained sequences with >70% coverage and >50% identity to establish CA homology. This search identified 11 belpaoviruses, all from nematode species, with amino-acid sequences and predicted structures closely related to those of Atlas virus (**Fig. 1d** and **Supplementary Table 1**). In addition to encoding phlebovirus-like Envs, the Gag proteins of this newly identified clade of Atlas-like viruses share belpaovirus hallmarks, including the absence of the N-myristylation motif in MA and the MHR in CA, and the presence of three zinc knuckles in NC. AlphaFold 3 models of their CA core structures superimposed closely, with a root mean square deviation of less than 1 Å, indicating a conserved fold (**Supplementary Table 1**).

### Recombinant Atlas-clade CA proteins self-assemble into icosahedral particles

To test whether CA domains from Atlas-clade viruses self-assemble, we expressed recombinant proteins comprising the CA core and flanking unstructured linkers from Atlas virus (GenBank: EPB78661) and two additional clade members, which we refer to henceforth as Atlas-Oos1 (GenBank: KAK6016282 from *Ostertagia ostertagi,* 83% identity to Atlas CA core) and Atlas-Nbr (GenBank: VDL73942 from *Nippostrongylus brasiliensis*, 65% identity to Atlas CA core; **Supplementary Table 1**). All three recombinant CA proteins were purified from *Escherichia coli* and spontaneously assembled into icosahedral capsid particles, most efficiently in high ionic strength phosphate buffers (**Fig. 2a** and **Supplementary Fig. 1**). Atlas CA showed the strongest self-assembly propensity, producing the highest particle-to-monomer ratio in size-exclusion chromatography (**Supplementary Fig. 1a**).

**Figure 2.**
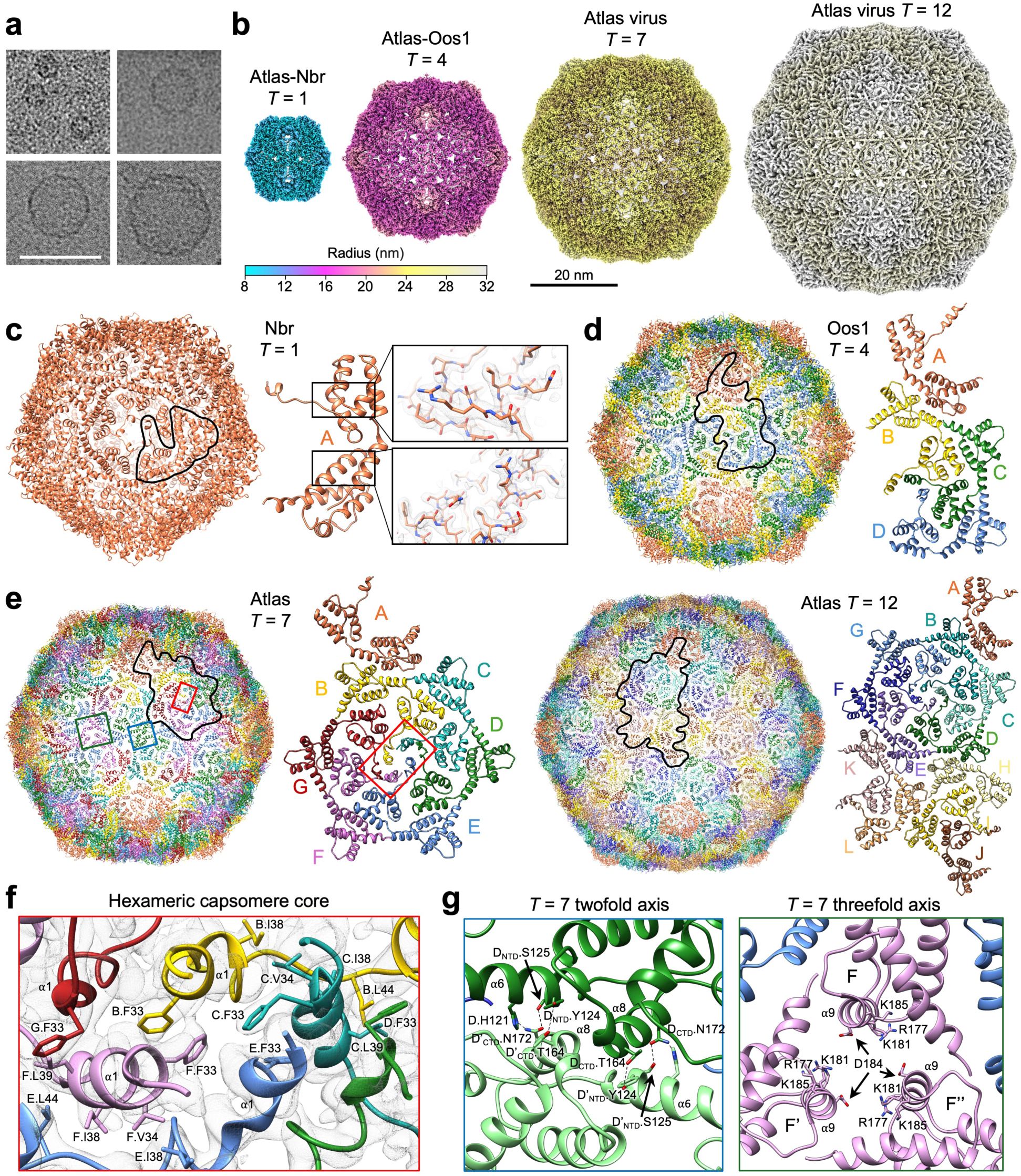
Atlas capsids assemble into icosahedral particles of different sizes. **(a)** Cryo-EM micrographs of four types of *in vitro*-assembled capsid particles from the three different nematode species. Top left, Atlas-Nbr (GenBank: VDL73942); top right, Atlas-Oos1 (GenBank: KAK6016282); bottom left and right, Atlas virus (GenBank: EPB78661). Scale bar, 50 nm. **(b)** Cryo-EM image reconstructions of Atlas-clade capsid particles, drawn to scale and colored by radius according to the color scale on the right with ChimeraX^59^. Scale bar, 20 nm. **(c)** Left, atomic model of the *T* = 1 Atlas-Nbr capsid with the asymmetric unit (ASU) outlined in black. Center, the ASU in cartoon representation. Right (insets), model with 2.2-Å resolution cryo-EM map. **(d)** Left, atomic model of the *T* = 4 Atlas-Oos1 capsid with the ASU outlined. Right, the ASU (chains A-D) in cartoon representation. **(e)** Atomic models of the *T* = 7 and *T* = 12 Atlas virus capsids. The ASUs are outlined and shown in cartoon representation as above. **(f)** Hydrophobic interactions and 3.2-Å resolution cryo-EM density at the center of a hexameric capsomere in the *T* = 7 Atlas virus particle (closeup of area in red box in panel (*e*)). **(g)** Capsid packing interactions in *T* = 7 Atlas virus particles at the twofold and threefold icosahedral symmetry axes (blue and green boxes in (*e*), respectively). See also Supplementary Fig. 3a-e.

Single-particle cryo-EM image reconstructions of the Atlas-clade CA particles yielded four structures at resolutions of 2.2-4.4 Å, sufficient for atomic model building and refinement (**Fig. 2b**, **Supplementary Table 2**). The linkers flanking CA (residues 1–28 and 214–244 of the Atlas CA expression construct, corresponding to polyprotein residues 160-187 and 373-403) were not resolved in any of the cryo-EM maps as predicted by AlphaFold 3. Deletion of Atlas CA residues 1-28 did not affect particle assembly. However, deletion of residues 214-244 reduced assembly efficiency and produced particles with imperfect icosahedral symmetry (**Supplementary Fig. 1e,f**), suggesting that the positively charged C-terminal tail of CA promotes efficient icosahedral capsid assembly.

All four particle structures displayed icosahedral symmetry but had different triangulation *(T)* numbers. Atlas-Nbr CA assembled exclusively into *T* = 1 particles with 60 protomers arranged as 12 fivefold-symmetric pentamers, yielding a 2.2 Å resolution structure (**Fig. 2c**). The other structures also contained 12 fivefold-symmetric pentamers along with different numbers and types of hexamers. Atlas-Oos1 CA predominantly formed *T* = 4 particles with 30 twofold-symmetric hexamers and a total of 240 protomers, yielding a 3.5 Å resolution structure (**Fig. 2d**). Atlas virus CA formed both *T* = 7 (420 protomers) and *T* = 12 particles (720 protomers), yielding reconstructions at 3.2 and 4.4 Å resolution, respectively (**Fig. 2e**). The *T* = 7 particles contained 60 hexamers in a single asymmetric environment, whereas the *T* = 12 particles contained 110 hexamers in three distinct environments: asymmetric, twofold-symmetric, and threefold-symmetric.

Despite the variations in particle size (20 to 60 nm diameter) and *T* numbers, the structures of individual protomers within pentameric capsomeres, exclusively located around icosahedral fivefold symmetry axes, are essentially identical in the four cryo-EM structures (**Supplementary Fig. 2a**). Quasi-equivalence between the pentameric and hexameric protomers is achieved primarily through a rotation of CA_NTD_ relative to CA_CTD_ by up to 7° (**Fig. 3a**). CA_NTD_ and CA_CTD_ are structurally identical in all capsomeres except for the pore-lining helix, CA_NTD_ helix α1, which adopts substantially different conformations in pentamers and hexamers, with an RMSD(Cα) of 4.9 Å. Due to this difference, a central pore is present in pentamers but absent in hexamers. The CA pentamer pore constricts to approximately 10 Å, a diameter geometrically compatible with passage of nucleotide-sized molecules (**Supplementary Fig. 2b**). The pore has a positive electrostatic potential due to a conserved basic residue on the luminal side of the pore, Lys37 in Atlas-Nbr CA, and Lys41 in Atlas virus CA (**Supplementary Figs. 2b,c** and **3a**).

**Figure 3.**
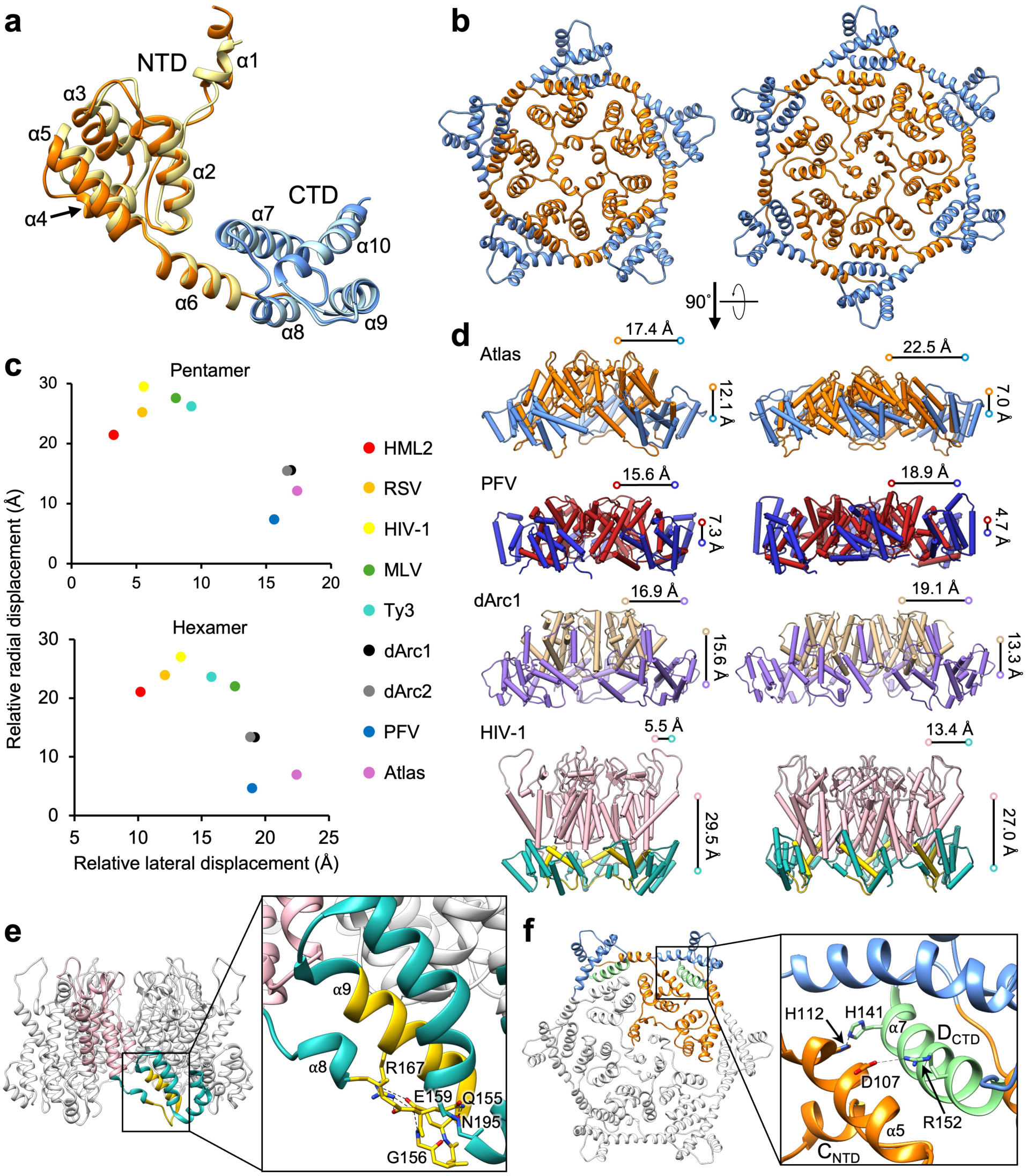
Atlas capsid assembly is reminiscent of spumaretroviruses and dArc. **(a)** Capsid (CA) protomer from a hexameric capsomere (darker colors, as in Fig. 1a,b) superimposed on a CA protomer from a pentameric capsomere (lighter colors) from the *T =* 7 Atlas virus particle. Orange, CA_NTD_. Blue, CA_CTD_. **(b)** Top views of pentameric and hexameric Atlas virus capsomeres. Orange, CA_NTD_. Blue, CA_CTD_. **(c)** Relative radial and lateral displacement of the centers of mass of CA_NTD_ and CA_CTD_ in retrovirus and retroelement capsomeres. HML2, human endogenous retrovirus-K HML-2 (betaretrovirus-like supergroup). RSV, Rous sarcoma virus (alpharetrovirus genus). PFV, prototype foamy virus (spumaretrovirus subfamily). dArc1/2, *Drosophila* dArc domesticated capsids. MLV, murine leukemia virus (gammaretrovirus genus). Ty3, retroelement capsid (metavirus genus). **(d)** Side views of pentameric and hexameric capsomeres from Atlas virus, PFV^33^, dArc1^13^, and HIV-1^47^. The HIV-1 major homology region (MHR) is highlighted in yellow. **(e)** Hydrogen bonding network formed by HIV-1 MHR. **(f)** Hydrogen bonding network formed by helix α7 of Atlas virus capsid, at the CA_NTD_-CA_CTD_ interface. Residue numbers refer to the CA expression construct (see Fig. 1a).

### Capsid assembly interactions in Atlas viruses

The pentameric capsomeres share a conserved architecture around the fivefold icosahedral symmetry axes in the Atlas-clade capsid structures. The hexameric capsomeres are more variable depending on whether or not they are centered on a two- or threefold symmetry axis. Overall, protomer packing is looser in pentameric capsomeres than in hexameric capsomeres. The average inter-protomer interface area in the Atlas virus lattice is 988 Å^2^ in pentamers versus 1186-1257 Å^2^ in hexamers. Despite differences due to location in the icosahedral lattice, all Atlas-clade capsomeres contain a conserved set of hydrophobic intra-capsomere contacts formed by the hydrophobic face of CA_NTD_ helix α1 of each protomer.

In asymmetric hexamers, found in the *T* = 7 and *T* = 12 particles, two pairs of opposing protomers, chains B-F and C-E, interact via π–π stacking between Phe33 side chains, thereby establishing a pseudo-twofold symmetry within the hexamers (**Fig. 2f**). Chains D and G adopt a different conformation, in which Phe33 instead packs against Leu39 of an adjacent protomer, either chain F or C, extending the hydrophobic network. Similar interactions are observed in the twofold-symmetric hexamers in the *T* = 4 and *T* = 12 particles, except that the opposing protomers are related by precise twofold symmetry. The α1 residues involved in these hydrophobic inter-protomer contacts (Phe33, Val34, Ala36, Ile38, Leu39, and Leu44 in Atlas virus CA) are conserved in Atlas, Atlas-Oos1, and Atlas-Nbr (**Supplementary Fig. 3a,e**). In more divergent clade members, conservative substitutions, including Phe33Tyr, Val34Phe, Ile38Leu, maintain the region’s hydrophobic character (**Supplementary Fig. 3a**). The conservation of these residues suggests they are important for capsid assembly. The presence of extensive hydrophobic inter-protomer interactions is consistent with the observed sensitivity of capsid assembly to low-salt conditions.

The *T* = 12 particles also contain hexamers with threefold symmetry (**Supplementary Fig. 4**), which precludes the pairwise Phe33-Phe33 π-π stacking observed in asymmetric and twofold-symmetric hexamers. Instead, the threefold-symmetric hexamers have a two-layered hydrophobic core formed by alternative packing of Phe33, Ala36 and Leu39, although we note that the side chain conformations remain uncertain due to relatively low (4.4 Å) resolution of the *T* = 12 structure (**Supplementary Fig. 4e**). Additionally, the threefold hexamers are less curved than the asymmetric and twofold hexamers (**Supplementary Fig. 4b-d**).

The strongest inter-capsomere contacts are a network of hydrogen bonds between CA_NTD_ helix α6 and CA_CTD_ helix α8 of the adjacent protomer. These contacts span the twofold axes in the *T* = 1 and *T* = 7 particles and pseudo-twofold axes in the *T* = 4 and *T* = 12 particles. In Atlas virus CA, the key contacts involve His121 and Ser125 to Asn172; and Tyr124 to Thr164 (**Fig. 2g**). Mutagenesis of His121 and Asn172 abolished capsid assembly, confirming the importance of these residues for capsid assembly (**Supplementary Fig. 1b**). His121 and Asn172 are conserved in all Atlas-clade members. The polar character of Ser125 is partially preserved as a threonine in some members (**Supplementary Fig. 3b-f**). The Tyr124-Thr164 pair is restricted to closer relatives of Atlas virus (Atlas-Oos1 and Atlas-Oos2), whereas position 164 exhibits significant polymorphism in more divergent members (**Supplementary Fig. 3b**). Additional inter-capsomere contacts occur at the threefold and pseudo-threefold axes in the *T* = 1, 4 and 7 assemblies. A basic patch on CA_CTD_ α9 interacts with a conserved aspartate, Asp184 in Atlas virus CA, in a neighboring protomer (**Fig. 2g** and **Supplementary Fig. 3e**). These electrostatic contacts are relatively long-range, typically exceeding 4 Å, and therefore probably provide weaker electrostatic stabilization at the inter-capsomere interface rather than salt bridges.

### Atlas capsid assemblies resemble spumaretroviruses and dArc

A prominent feature of Atlas capsid assemblies is their thin shell, which differs from the more radially stratified organization typical of many retrovirus and retroelement capsids. In canonical retroviral assemblies, CA_NTD_ and CA_CTD_ are arranged in two radial layers, with CA_NTD_ forming the outer layer and the CA_CTD_ forming the inner layer of the protein shell. By contrast, in the Atlas capsids, particularly in hexameric capsomeres, the centers of mass of CA_NTD_ and CA_CTD_ have greater lateral than radial displacement (**Fig. 3b,c**). This produces a thin shell in which CA_NTD_ occupies the capsomere center and CA_CTD_ decorates the capsomere periphery (**Fig. 3b,d**).

This geometry is most reminiscent of prototype foamy virus (PFV), a spumaretrovirus that forms large, thin-shelled *T* = 13 capsids (**Fig. 3d**)^33^. To a lesser extent, Atlas particles also resemble the dArc1 and dArc2 assemblies from *Drosophila* species, which form *T* = 4 particles with a similarly less stratified CA_NTD_ and CA_CTD_ arrangement (**Fig. 3c,d**)^13^. These similarities appear to reflect convergent or analogous lattice organization rather than genetic conservation: Atlas, spumaretrovirus and dArc capsid proteins share no significant amino acid sequence identity, and their principal capsid assembly interfaces are not conserved.

Atlas and PFV both lack the MHR and an N-myristoylation motif, a combination rarely observed in retroviral Gag proteins. *Drosophila* dArc capsids contain a putative MHR-like sequence with limited similarity to the orthoretrovirus MHR. The canonical orthoretrovirus MHR is a 20-residue motif in the N-terminal region of CA_CTD_ that forms an intrachain hydrogen-bonding network, which structurally underpins CA_CTD_ (**Figs. 1d, 3e**)^41^. In HIV-1, Lys158 in the MHR forms a positively charged ring in the central pore of immature hexameric capsomeres that recruits inositol hexaphosphate (IP_6_), a cofactor essential for lattice stabilization and capsid maturation^42–44^. Mutations within the MHR impair retroviral capsid assembly^45^.

Lacking an MHR, the Atlas viruses contain an alternative set of polar intra-capsomere contacts, between CA_NTD_ helix α5 and CA_CTD_ helix α7 of a neighboring protomer, that drives capsid assembly. The most notable of these contacts is a conserved salt bridge, formed by Asp107 and Arg152. This is supplemented in Atlas virus and Atlas-Oos1 by a hydrogen bond between His112 and His141 (**Fig. 3f**; **Supplementary Fig. 3c**; residue numbers refer to Atlas virus CA). Mutation of His112 abolished Atlas virus capsid assembly. Because Atlas hexamers deviate from perfect sixfold symmetry, these CA_NTD_-CA_CTD_ contacts are only formed at two chain interfaces in asymmetric and twofold-symmetric hexamers (involving chains C-D, F-G, or H-I and their symmetry-related copies), and at three interfaces in threefold-symmetric hexamers. In the more loosely packed pentameric capsomeres, the Asp107-Arg152 and His112-His141 residue pairs are too far apart (>5 Å) to form stable interactions.

### Comparison of Atlas and HIV-1 capsomere pore structures

Atlas capsid pentamer pores are lined with a ring of basic residues 15 Å in diameter, formed by Lys41 in Atlas virus and Lys37 in Atlas-Nbr (**Supplementary Figs. 2b and 5**). The pore geometry and positive electrostatic charge are reminiscent of HIV-1 capsomere pores, in which rings of basic residues coordinate IP_6_. In the immature HIV-1 capsid hexamers, Lys158 in the MHR contributes to the IP_6_ binding site, along with Lys227 from spacer peptide SP1^44^. In mature HIV-1 capsid pentamers and hexamers, IP_6_ is coordinated by Arg18 and Lys25 in CA_NTD_, forming two stacked electropositive rings in the capsomere pore (**Supplementary Fig. 5**)^42,43^. This raised the question of whether IP_6_ binds to Atlas capsid pentamers and hence contributes to Atlas capsid assembly.

Addition of IP_6_ did not alter *in vitro* Atlas virus capsid morphology or assembly efficiency (**Supplementary Fig. 5b**). Consistent with this, HIV-1 capsomeres have two rings of basic residues to bind IP_6_, formed by Arg18/Lys25 in mature capsids or Lys158/Lys227 in immature capsids. Atlas virus pentamer pores contain a single basic residue ring. Moreover, the 15-Å ring diameter in Atlas pentamers is larger than the 8-9 Å diameter of the rings with high IP_6_ binding affinity in HIV-1, those formed by Arg18, Lys158 and Lys227, and instead resembles the low-affinity, auxiliary IP_6_-binding ring formed by Lys25 (**Supplementary Fig. 5c**)^42–47^. In Atlas capsid hexamers, the basic residues do not form rings, due to the deviation of the hexamers from sixfold symmetry, precluding IP_6_ coordination. Together, available structural comparisons and biochemical data suggest that although inositol phosphate coordination by Atlas pentamers is geometrically possible, the number and spacing of pore-lining basic residues differ substantially from those of the high-affinity IP_6_-binding sites in HIV-1. We conclude that inositol phosphates are unlikely to function as pocket factors or cofactors in Atlas capsid assembly.

### Atlas particles package nucleic acids

Atlas virus capsid particles copurified with nucleic acids based on their UV-range absorbance spectrum, with a 260:280 nm absorbance ratio consistently above 1 and approaching 2 in many preparations. The copurified nucleic acids resisted repeated nuclease treatment and high-salt washes during purification (**Fig. 4a**). Cryo-EM reconstructions did not reveal ordered nucleic acid density, possibly due to icosahedral symmetry averaging. However, in a subset of particles dense material was visible near the inner capsid surface, consistent with packaging of nucleic acids from the bacterial cells used for capsid expression (**Fig. 4b**). Similar density features were observed when capsids were assembled in the presence of 5 kb plasmid DNA and subsequently treated with nuclease (**Fig. 4b**).

**Figure 4.**
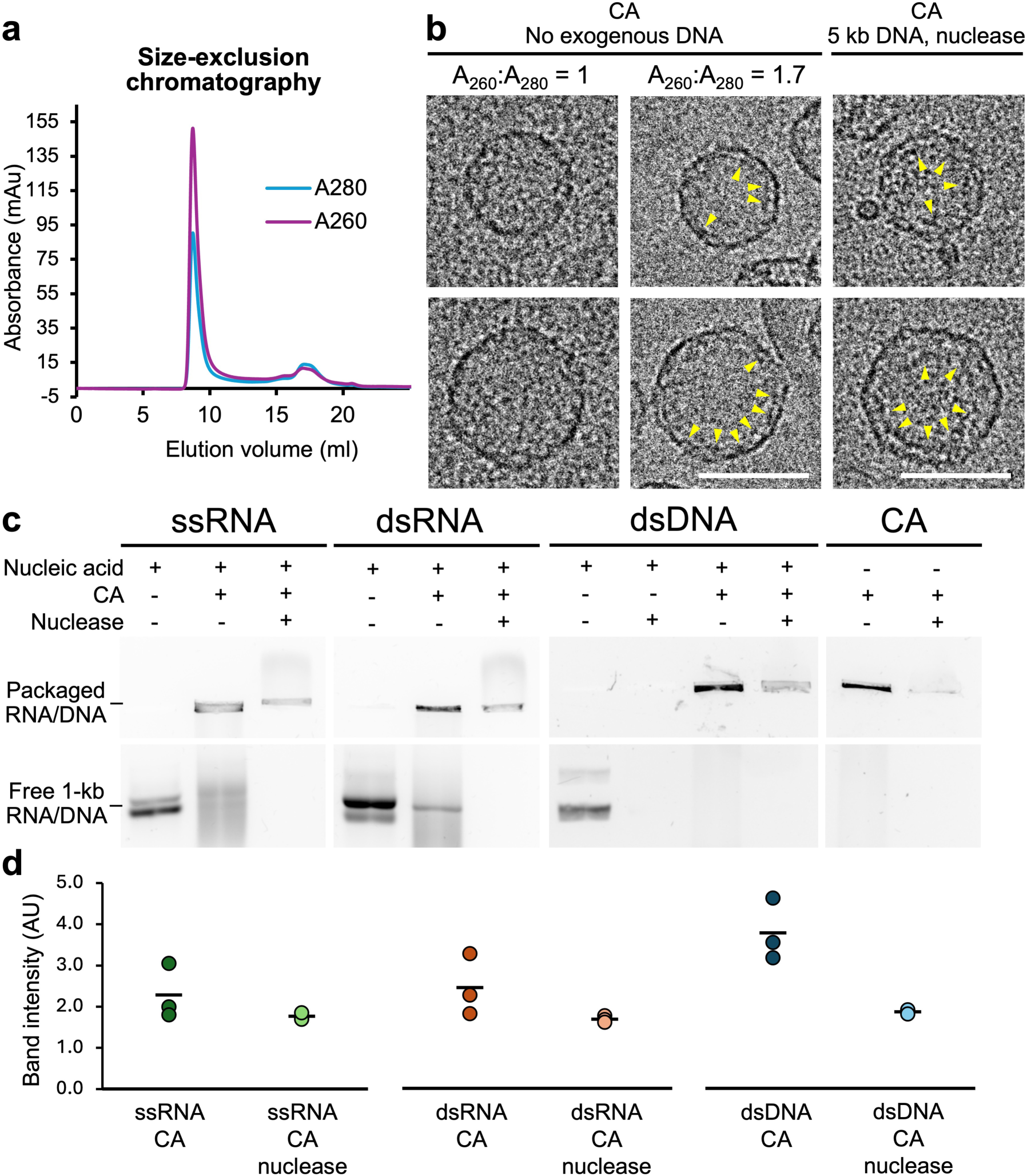
Intrinsic nucleic acid packaging activity of Atlas capsid particles. **(a)** Size-exclusion chromatography traces of Atlas virus capsids. The high 260:280 nm absorbance ratio (A_260_:A_280_) indicates that nucleic acids were copurified with the capsids. **(b)** Cryo-EM micrographs of Atlas virus *T* = 7 and *T* = 12 capsids assembled in the absence of exogenous nucleic acids (left and center panels); or in the presence of 5 kb plasmid DNA followed by salt active nuclease treatment (right). Yellow triangles indicate density attributable to nucleic acids. Nucleic acid densities were more visible in samples with higher A_260_:A_280_ or with exogenous nucleic acids. Scale bars, 50 nm. **(c)** Electrophoretic mobility shift assay (EMSA) of Atlas virus capsids (CA) with or without application of the indicated extrinsic nucleic acid; and with or without treatment of Benzonase nuclease. **(d)** Quantification of nucleic acid protection, measured as band intensity of packaged nucleic acid. Data from three independent experiments are shown with a line showing the mean.

To assess nucleic acid packaging by Atlas capsids we performed electrophoretic mobility shift assays (EMSAs). Bands corresponding to free 1-kb ssRNA, dsRNA, or dsDNA were depleted when Atlas capsid protein was present, indicating that all three nucleic acid types bound to the capsid (**Fig. 4c**). Moreover, capsid-bound nucleic acids were partially nuclease-resistant, consistent with packaging, or encapsidation: nuclease treatment eliminated free nucleic acids not particle-associated nucleic acids. EMSA quantification showed retention of 77% ssRNA, 69% dsRNA, and 49% dsDNA (**Fig. 4d**).

The lack of selectivity for nucleic acid type or sequence suggests encapsidation occurs via nonspecific electrostatic interactions. Atlas capsid is highly basic, with a predicted pI of 9.2 and 28 basic residues, most of which lie in the folded CA core and map to the inner capsid surface. Additionally, five of the basic residues are located in the unresolved 31-residue C-terminal tail (pI = 11.0), also located in the capsid lumen. The capsid lumen therefore has a strong positive electrostatic potential and could plausibly bind nucleic acids without the NC zinc knuckles, which were absent in the capsid expression construct (**Supplementary Fig. 6a**). Deletion of the C-terminal tail reduced the yield and 260:280 nm absorbance ratio of purified capsid particles, indicating reduced nucleic acid packaging (**Supplementary Fig. 1f**). Thus, the C-terminal tail may promote capsid assembly by nucleating oligomerization on nucleic acid, analogous to NC in retroviral assembly^40^.

Atlas-Oos1 and Atlas-Nbr capsid assemblies also copurified with nucleic acids, albeit with lower 260:280 nm absorbance ratios than Atlas (**Supplementary Fig. 1c,d**). The Atlas-Oos1 CA core has a net negative charge (**Supplementary Fig. 6b**), and its nucleic acid packaging activity may depend mainly on its basic unstructured C-terminal tail. Atlas-Nbr has a similar luminal charge distribution to Atlas virus; the lower nucleic acid packaging of *T* = 1 Atlas-Nbr particles may instead be due to their much smaller internal volume.

### Atlas particles are spontaneously endocytosed and disassemble at endosomal pH

Having established that Atlas particles can encapsidate RNA and DNA, we sought to determine whether the particles could enter cells. We tracked Atlas particles covalently labeled with Alexa Fluor 647 by fluorescence microscopy in HEK293T, HeLa and mouse iBMDM cells. Within 30 minutes, labeled particles accumulated in perinuclear puncta in all three cell types (**Fig. 5a,b**). Colocalization with LysoTracker dye indicated that the internalized protein accumulated in endolysosomal compartments (**Fig. 5b**). Atlas-Oos1 and Atlas-Nbr particles showed similar entry patterns (**Fig. 5c**). Uptake was temperature dependent – endosomal CA puncta appeared by 30 minutes and increased up to 90 min at 37 °C (**Fig. 5c**) but no CA puncta were observed at 4 °C, confirming that cell entry occurred by active endocytosis.

**Figure 5.**
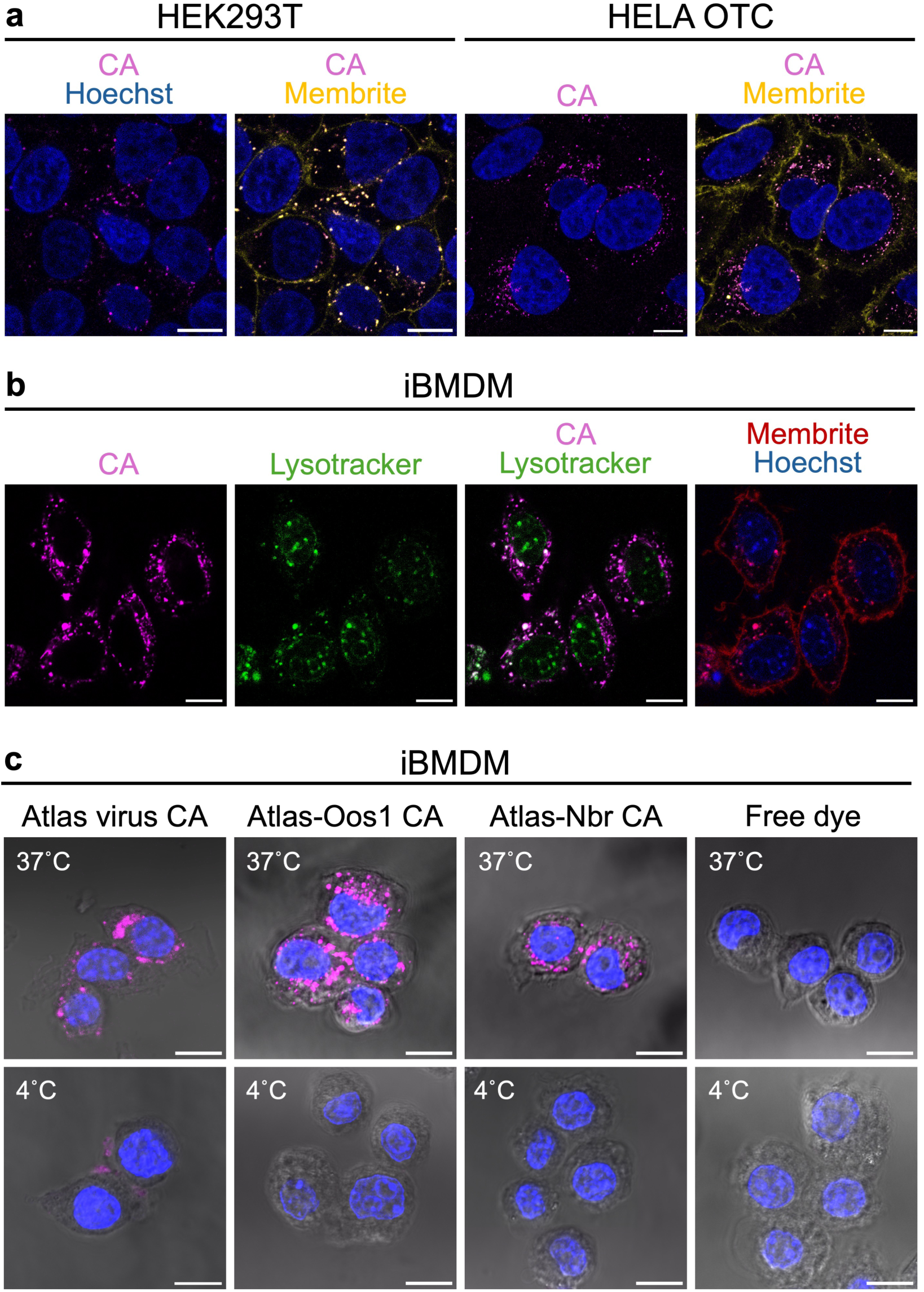
Endocytic cell entry of Atlas capsid particles. **(a)** Live cell imaging of Atlas virus capsid uptake by human HEK293T and HeLa cells. Magenta, fluorescently labeled capsids. Blue, Hoechst nuclear stain. Yellow, MemBrite plasma membrane dye. Scale bars, 10 µm. **(b)** Live cell imaging of Atlas virus capsid uptake by mouse immortalized bone marrow-derived macrophages (iBMDMs). Magenta, fluorescently labeled capsids. Green, LysoTracker Green endolysosome dye. Red, MemBrite plasma membrane dye. Blue, Hoechst nuclear stain. Scale bars, 10 µm. **(c)** Fixed cell imaging of fluorescently labeled Atlas virus, Atlas-Oos1, and Atlas-Nbr capsid uptake by iBMDMs at 37°C and 4°C, with free dye control (unconjugated Alexa Fluor 647). Magenta: fluorescently labeled capsids (Alexa Fluor 647). Blue, Hoechst nuclear stain. Scale bars, 10 µm.

Since Atlas particles accumulate in endolysosomes, we next assessed their stability under acidic conditions. At pH 4.5, Atlas virus particles dissociated and released encapsulated nucleic acids (**Fig. 6**). This pH sensitivity is likely attributable to histidine residues located at inter-protomer interfaces, including His112, His121 and His141 in Atlas virus CA. Protonation of histidine side chains at endolysosomal pH is expected to weaken hydrogen bonds and compromise particle integrity. As noted above, mutation His112 or His121 abolished particle assembly. In summary, our data show that Atlas particles package kilobase-scale RNA and DNA, are endocytosed, accumulate in acidic endolysosomal compartments, and disassemble at low pH. We note that these are all desirable properties for nucleic-acid delivery vehicles.

**Figure 6.**
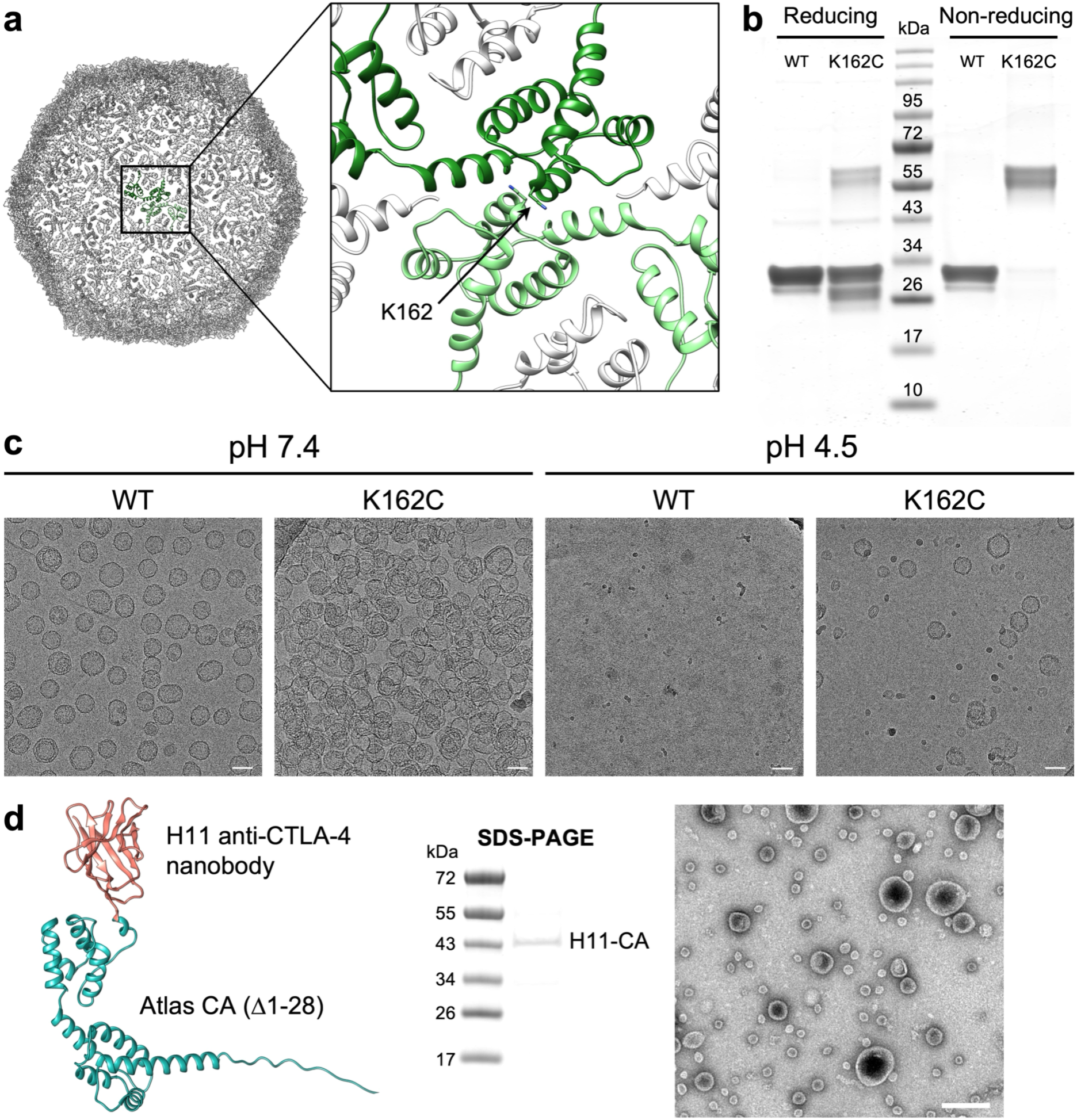
Engineering the capsid particles as nucleic acid delivery tools. **(a)** Lysine 162 is located near a symmetry related copy at the icosahedral twofold symmetry axis (shown here) and pseudo-twofold axes in Atlas virus capsid particles. **(b)** SDS-PAGE of WT and K162C Atlas virus capsid in reducing and non-reducing conditions. K162C capsids form disulfide-linked dimers. **(c)** Cryo-EM micrographs of WT and K162C capsids at pH 7.4 or pH 4.5. Scale bars, 50 nm. **(d)** Left: AlphaFold 3 model of the anti-CTLA-4 nanobody H11^48^ fused to the N-terminus of Atlas virus CA. Center: SDS-PAGE of the purified H11-CA fusion protein. Right: negative stain electron micrographs of H11-CA particles. Scale bar, 100 nm.

### Engineering Atlas capsids for pH resilience and receptor-directed ligand display

Having obtained Atlas particle structures, we sought to demonstrate their usefulness in structure-guided design. First, we engineered an intermolecular disulfide at the icosahedral twofold to increase particle stability. Lys162, near the N-terminus of helix α8, is located near its symmetry- or pseudosymmetry-related copies in Atlas particles (**Fig. 6a**). The K162C substitution resulted in particles containing disulfide-linked CA dimers throughout the lattice (**Fig. 6b**). K162C particles showed increased particle clustering at neutral pH, but a substantial fraction remained intact after acidification of the buffer to pH 4.5, whereas WT capsids fully dissociated (**Fig. 6c**). Under the same conditions, a subset of K162C particles formed smaller spherical assemblies (**Fig. 6c**). We infer that covalent stabilization of CA-CA interfaces enhances pH tolerance and provides a route for increasing particle stability in applications where cargo protection is desirable.

Next, we aimed to engineer Atlas capsids to display a specific receptor ligand. Some nucleic acid-based therapies require cell-type targeting, but this remains a challenge for current delivery platforms^22^. The Atlas capsid N-terminus is exposed on the exterior particle surface, providing a site for modular functionalization. We replaced the unstructured N-terminal tail of Atlas CA, which is dispensable for assembly, with the H11 anti-CTLA-4 nanobody^48^. The T cell-associated CTLA-4 receptor is a major target in checkpoint blockade cancer immunotherapy^48^. The H11-CA fusion protein retained the ability to assemble into particles (**Fig. 6d**). Together, our data provide proof-of-principle for two complementary strategies to engineer Atlas capsids, covalent stabilization and receptor-binding ligand display.

## DISCUSSION

Here, we identify the Atlas viruses as a clade of eleven endogenous viruses in nematodes with distinctive genetic and structural features. Members of the *Belpaoviridae* family^30^, Atlas viruses have retrovirus-like Gag-Pol-Env genomic organization but their Env contains a phlebovirus-like class II membrane fusogen not previously reported in reverse-transcribing viruses^29^. Atlas Gag proteins also differ substantially from orthoretrovirus Gags: they share no significant sequence homology and lack the major homology region (MHR) and N-myristoylation signal required for orthoretrovirus capsid assembly and membrane targeting, respectively. Our cryo-EM data reveal that the capsids from three representative Atlas virus species form thin-shelled icosahedral assemblies of distinct outer diameters ranging from 20 to 60 nm. These assemblies and their constituent capsomeres are structurally distinct from those of orthoretrovirus capsids but reminiscent of spumaretrovirus (foamy virus) capsids^33^ and to a lesser extent *Drosophila* dArc capsids^13^. In Atlas and foamy virus capsomeres CA_CTD_ is located on the side of CA_NTD_ distal to the capsomere center rather than on the side of CA_NTD_ facing the particle lumen. Hence, both domains contribute to the outer particle surface rather than forming two stratified layers as in orthoretroviruses (**Fig. 3, Supplementary Fig. 5**). These findings expand the known genetic and structural diversity of viral structural proteins that has accompanied host-virus coevolution.

Like Atlas virus Gags, foamy virus Gags lack an N-myristoyl group and are therefore not targeted to the plasma membrane. Instead, foamy virus capsids assemble intracellularly and particle egress requires specific Gag-Env interactions^37^. Foamy virus Envs are also structurally divergent from orthoretrovirus Envs and instead resemble the F proteins of paramyxo- and pneumoviruses^33,49^. The presence of a functional Env and self-assembling Gag raises the possibility that Atlas-clade members may retain the capacity to form enveloped virions. Further studies are required to determine whether this is the case, ideally by expressing the full Gag-Pol-Env sequence in relevant cellular contexts.

Beyond their evolutionary implications, Atlas capsid particles have properties relevant to nucleic acid delivery. Their intrinsic properties include packaging of kilobase-scale RNA and DNA without sequence specificity, endocytic cell-entry, accumulation in acidic endosomal compartments, and low pH-triggered disassembly. The structures reported here provide a framework for engineering new functionality into these particles. As a proof-of-concept, we successfully designed particles with increased pH resilience by introducing a disulfide bond at the capsid lattice interface, and we generated particles displaying a CTLA-4 T cell-targeting tag. These results establish Atlas capsids as a genetically programmable protein nanoparticle scaffold.

Recombinant Atlas capsid particles have potential advantages over existing virus-based nucleic acid delivery systems. With internal diameters of up to 55 nm for the T = 12 Atlas particles, their internal volumes are sufficient, in principle, to accommodate nucleic-acid cargos larger than those packaged by AAVs^24^, including payloads for CRISPR-Cas9 gene editing^27^ and emerging *in vivo* CAR-T approaches^28^. Consistent with this, the complete Atlas virus genome spans 9.2 kb^29^. The broad size range of Atlas particles may also provide opportunities for cargo-matched particle selection, with smaller particles potentially advantageous where particle size limits tissue distribution. As small recombinant proteins, Atlas capsids can readily be produced at scale. Lacking an integrase and polymerase, Atlas particles will bypass the risks of off-target integration associated with retroviral and lentiviral vectors^20,21,25^. Human retroelement-derived capsids are currently being explored as RNA delivery vehicles^20,21^, but their endogenous physiological roles warrant careful safety evaluation in ectopic or high-dose use. Although their immunogenicity remains to be tested, the nematode origin of Atlas capsids limits overlap with human retroelement biology and reduces the likelihood of preexisting immunity compared with human-tropic viral scaffolds. Together, the Atlas capsid structures provide a rational foundation for developing programmable protein-based nanoparticles for nucleic-acid delivery. Future studies are needed to convert these structural and biochemical properties into functional delivery performance, including determining whether Atlas capsid functionalization can be used more broadly to achieve cell type-specific payload delivery, validating cargo activity after uptake, improving endosomal escape, testing tissue-selective targeting and evaluating safety after repeated or high-dose administration.

## METHODS

### Phylogenetic analysis

Atlas virus (GenBank: EPB78661 from *A. ceylanicum*) was originally identified via PSI-BLAST searches for phlebovirus-like proteins^29^. Its G_C_-like Env was the closest match among sequences outside known infectious virus taxa. To delineate MA, CA, and NC, we aligned the Atlas Gag sequence to well-characterized retroviral Gag proteins (HIV-1 and MLV) using Clustal Omega^50^ and predicted the Atlas Gag structure with AlphaFold 3^34^. We defined the Atlas clade with a two-step BLASTP pipeline: (i) PSI-BLAST query of nematode genomic databases with the entire 2,828 a.a. Atlas virus sequence, retaining hits with >70% coverage; (ii) BLASTp query of the filtered ERV-like hits using the structured region of Atlas CA (a.a. 188–372), retaining hits with >70% coverage and >50% identity. Eleven belpaoviruses met these criteria and were classified within the clade. Phylogenetic relationships were generated with iTOL v7^51^. Sequence alignments and logos were generated with ESPript3.2^52^ and WebLogo3^53^, respectively (**Supplementary Fig. 3**).

### Protein expression

Genes encoding the CA domain of Atlas virus (residues 160-403 of the polyprotein), Atlas-Oos1 (GenBank: KAK6016282 from *O. ostertagi*), and Atlas-Nbr (GenBank: VDL73942 from *N. brasiliensis*) were codon-optimized for expression in *E. coli* and cloned into vector pGEX-6P-1 in frame with the GST tag and the HRV 3C site by blunt end Gibson assembly. A 6 amino-acid linker containing triple Gly-Ala repeats was added between the HRV 3C site and the gene insert. *A. ceylanicum* CA mutants were generated by the Q5 site-directed mutagenesis kit (NEB). The fusion construct was generated by replacing the *A. ceylanicum* MA-CA disordered loop with H11 nanobody and cloning into vector pET-49b in frame with the GST tag and His tag.

Chemically competent *E. coli* BL21(DE3) cells were transformed with the plasmids described above and plated on LB agar containing selection antibiotics for overnight incubation at 37 °C. Individual colonies were picked and grown in LB supplemented with antibiotics overnight at 37 °C, 180 RPM. The starter cultures were used to inoculate expression cultures at a 1:100 ratio. Expression cultures were grown at 37 °C, 180 RPM until they reached the exponential growth phase with OD600 = 0.6-0.8. Protein expression was induced by adding 400 µM IPTG and grown overnight with ampicillin or kanamycin at 18 °C, 180 RPM.

### Protein purification

The affinity purification protocol was modified from Pastuzyn *et al.*^12^. *E. coli* expression cultures containing the pGEX-6P-1-CA plasmid were grown overnight at 18 °C, 180 RPM in the presence of 400 µM IPTG. Cells were harvested by centrifugation at 5000 g for 15 min at 4 °C. For every 500 ml of culture, the pellet was resuspended in 10 ml lysis buffer (500 mM NaCl, 50 mM Tris-HCl, pH 8.0, 5% glycerol) and flash frozen in liquid nitrogen. Frozen pellets were thawed and supplemented with lysozyme; salt active nuclease (Sigma-Aldrich); cOmplete EDTA-free protease inhibitor cocktail (Sigma-Aldrich); DTT, to a concentration of 1 mM. Cells were sonicated for 8 min with 5 s pulses, 33% duty cycle, and centrifuged at 30,000 g for 45 min at 4 °C. Supernatant was passed through a 0.45-µm filter and incubated with pre-equilibrated Pierce glutathione agarose resin (Thermo Fisher Scientific) in a gravity flow column for 90 min at 4 °C on a shaker. The bound protein was washed with 25 resin bed volumes of lysis buffer and cleaved with PreScission Protease (GE Healthcare) in cleavage buffer (150 mM NaCl, 50 mM Tris-HCl pH 7.2, 1 mM EDTA, and 1 mM DTT) overnight at 4 °C. Untagged CA was directly eluted, and the bound GST-protease complex was eluted by 20 mM reduced L-glutathione, 10 mM Tris-HCl pH 7.4.

Purified CA was dialyzed overnight at 4 °C against assembly buffer (1 M NaCl, 2×PBS, 1 mM DTT), with or without 1 mM IP_6_ pH 7.4. Following buffer exchange, CA was concentrated and incubated at 37 °C for 15 min, then chilled at 4 °C for 45 min before being loaded on a Superdex 200 Increase 10/300 GL column (Cytiva) for size-exclusion chromatography. Elution fractions were mixed with reducing or non-reducing Laemmli SDS sample buffer (Invitrogen) and analyzed by electrophoresis with NuPAGE Bis-Tris gels (Invitrogen).

For the purpose of visualizing nucleic acid packaging by cryo-EM, capsids were prepared as described above, with or without addition of purified 5 kb plasmid DNA at a final concentration of 1/5^th^ of the capsid particle concentration. For DNA-supplemented samples, Salt Active Nuclease (New England Biolabs) was added 10 min prior to vitrification.

### Negative stain EM

Fractions from the void elution peak of size-exclusion chromatography were collected and diluted to 0.1 mg/ml. Carbon film-coated copper 200-mesh grids (Agar Scientific) were glow discharged for 90 s in a vacuum chamber at 30 mA. 2 µl of sample was incubated on the grid for 1 min. Excess sample was removed by side-blotting with a filter paper. 2 µl 2% uranyl acetate was applied to the grid twice with immediate side-blotting, and a third time with 1 min incubation before blotting. Negative stain specimens were allowed to dry at RT for at least 12 h. Micrographs were taken by a 120 kV Tecnai Spirit microscope (Thermo Fisher Scientific) equipped with a Gatan Orius CCD camera.

### Cryo-EM

R1.2/1.3 or R2/2 300 mesh copper grids (Quantifoil Micro Tools, Germany) were glow discharged for 75 s at 30 mA. 3 µl 0.2 mg/ml graphene oxide (Sigma-Aldrich) was applied to the carbon side of glow discharged grids, incubated for 15 s, and side-blotted on filter paper. Grids were washed by adding and blotting 10 µl MilliQ water twice on the carbon side, and once on the metal side. Grids were stored at room temperature for at least 12 h to dry. Freezing was performed by a Vitrobot Mark IV (Thermo Fisher Scientific) at 4 °C and 100% relative humidity. 2 µl of 3 mg/ml sample was applied to the coated side of grids, blotted for 4-6 s at -5-5 force and plunge frozen. Cryo-EM data collection was performed on a 300 kV Titan Krios microscope (Thermo Fisher Scientific) equipped with a 20-eV slit-width Gatan BioQuantum energy filter and a Gatan K3 direct electron detector in counting mode. Movies were recorded by EPU (Thermo Fisher Scientific) at a magnification of 105,000× (pixel size 0.826 Å) and a dose rate of 1 e^-^ Å^-2^ s^-1^ per frame. For visualizing nucleic acid packaging (**Fig. 4b**), micrographs were acquired on a 200 kV Glacios microscope (Thermo Fisher Scientific) equipped with a Falcon III direct electron detector.

Movies were processed in cryoSPARC v4^54^. All movies were aligned by the Patch Motion Correction algorithm of cryoSPARC. The contrast transfer function parameters were estimated by Patch CTF. 1,000 particles of each size group were manually picked and 2D classified to generate a template for automated blob picking. Particles were extracted with a cubic box size of 450, 700, 1000, and 1200 pixels for *T* = 1, *T* = 4, *T* = 7, and *T* = 12 particles, respectively. For the *T* = 12 particles the box was Fourier-cropped to 600 pixels during refinement due to GPU memory constraints.

### Model building and refinement

AlphaFold models of the CA_NTD_ and CA_CTD_ served as starting coordinates. Each domain was rigid-body fitted into the reconstructed density in UCSF Chimera^55^ with the Fit-in-Map command, and unresolved N- and C-terminal loops were trimmed. The domains were sequentially docked and linked to compose one asymmetric subunit. The cryo-EM map was boxed around this model in PHENIX^56^ with the Map Box function, followed by Real Space Refinement with restraints on secondary structure, non-crystallographic symmetry between chains, sidechain rotamer, and Ramachandran. Manual rebuilding to resolve steric clashes and geometry outliers was performed in Coot^57^. Finally, icosahedral symmetry was expanded from the asymmetric subunit to produce the full-particle atomic model.

### Structure analysis

For structure comparison between Atlas clade members, the model alignment and RMSD calculation was done in UCSF Chimera using the MatchMaker function. The AlphaFold 3 model of each CA monomer was aligned with the structurally resolved Atlas CA monomer in its capsid pentamer. The resolved pentamers of Atlas, Atlas-Oos1 and Atlas-Nbr were aligned. A single chain conformation within Atlas pentamer and hexamer was aligned.

For comparison of retroviral and retrotransposon capsomere architecture, the method was adapted from Calcraft *et al.*^33^. The following PDB models were used: dArc1/dArc2 pentamer: 6TAR/6TAT^13^, hexamer: 6TAS/6TAU^13^; Ty3 pentamer and hexamer: 6R24; HML2 pentamer: 6SSJ, hexamer: 6SSM; HIV-1 pentamer: 5MCY^47^, hexamer: 5MCX^47^; MLV pentamer: 6HWY, hexamer: 6HWX; RSV pentamer: 7NO0, hexamer: 7NO5; PFV pentamer: 8OZL^33^, hexamer: 8OZM^33^. For each structure, CA_NTD_ and CA_CTD_ residue ranges were defined (**Supplementary Table 1**), and Cα coordinates were extracted. For each chain, Cα coordinates within the domain were averaged to yield a chain-level centroid, forming CA_NTD_ and CA_CTD_ rings. The oligomer axis was estimated as the unit normal to the best-fit plane through the ring centroids by principal-component analysis (PCA). Average ring radii (R_NTD_, R_CTD_) were computed as the mean distances of centroids to the axis; lateral displacement was ΔR = R_NTD_ − R_CTD_. Axial displacement was ΔZ = (mean_NTD_ − mean_CTD_)·n, i.e., the projection of the CA_NTD_-CA_CTD_ centroid offset onto the axis. Analyses were implemented in custom Chimera Python scripts operating on Cα atoms, with distances reported in Å. Electrostatic surfaces were calculated with APBS (https://server.poissonboltzmann.org)^58^ and displayed with 3DMol or ChimeraX^59^. See **Supplementary Table 2** for additional data collection parameters.

### Electrophoretic mobility shift assay (EMSA)

After protease cleavage, eluted tag-free Atlas virus CA was either left untreated or pre-incubated with 10 µg ssRNA, dsRNA, or dsDNA. Samples were concentrated at 4 °C using centrifugal filters (Amicon) and buffer-exchanged into 1× PBS (137 mM NaCl, 2.7 mM KCl, 4.3 mM Na_2_HPO_4_, 1.47 mM KH_2_PO_4_). Concentrated CA was loaded onto a Superdex 200 Increase 10/300 GL column (Cytiva), and void-volume fractions were collected. Each preparation was split equally: one aliquot was left untreated and the other treated with Benzonase (Sigma-Aldrich) for 10 min at 4 °C following the manufacturer’s instructions. Additional controls comprised free nucleic acids treated with Benzonase under the same conditions. Samples were mixed with NativePAGE sample buffer (Thermo Fisher Scientific) and run on 0.7% agarose gels stained with 1× SYBR Safe (Thermo Fisher Scientific) in 1× TBE (90 mM) for 60 min at 100 V at 4 °C. Gels were imaged on a G-BOX system (Syngene). Band intensities were measured in ImageJ/Fiji^60^.

### Cell entry assay

Purified capsid particles were fluorescently labelled with the Alexa Fluor 647 Antibody Labeling Kit (Invitrogen). The succinimidyl-ester dye was prepared in 50 µl assembly buffer (2× PBS, 1 M NaCl, 1 mM DTT), and 5 µL of the unconjugated dye was retained as a control. Protein (0.5 ml at 2 mg/ml) was supplemented with 0.1 M NaHCO₃ and incubated with dye for 1 h at room temperature. Excess dye was removed by centrifugation through Zeba dye and biotin removal spin columns (Thermo Fisher Scientific).

HEK293T, HeLa, and immortalized primary mouse bone marrow-derived macrophages (iBMDMs) were cultured in high-glucose DMEM (Gibco) supplemented with 10% (v/v) FBS (Gibco). 24 h before experiments, cells were seeded onto 35 mm µ-Dishes or µ-Slide 4-well chambers (ibiTreat, Ibidi) at 5×10⁴ cells/ml to form monolayers. For live-cell imaging, nuclei were stained with Hoechst 33342 (PureBlu Bio-Rad; 11 ng/ml, 15 min, 37 °C), acidic compartments with LysoTracker Green DND-26 (Cell Signaling Technology; 1 µM, 45 min, 37 °C), and the plasma membrane with MemBrite Fix (Bio-Rad; 10 min, room temperature). 20 µl of fluorescently labelled capsids or unconjugated dye were added to each µ-Dish and imaged immediately by confocal microscopy.

For fixed-cell imaging, 5 µl of labelled capsids or unconjugated dye were added to each well of the 4-well chamber. Cells were incubated at 37 °C or 4 °C for 30 or 90 min and fixed with 4% paraformaldehyde for 10 min at room temperature. Fixed cells were counterstained with Hoechst 33342 (PureBlu, 11 ng/ml, 10 min, room temperature) prior to imaging. The images were processed with ImageJ/Fiji^60^.

### Fluorescence confocal microscopy

Single point-scanning confocal microscopy was carried out on a Zeiss LSM 780 microscope using a 63×/1.4 NA Plan-apochromat oil immersion objective lens. The microscope was equipped with 405, 458, 488, 514, 561 and 633 nm laser lines. For live-cell imaging, excitation at 405, 488, 561, and 633 nm was used with emission collected at 410-484, 490-544, 552-605, and 641-756 nm, respectively. Time-lapse acquisition was conducted for 90 min at 37 °C in 5% CO₂ using an on-stage incubator. For fixed-cell imaging, the 633 nm line was used with emission collected at 641-756 nm.

### pH disassembly assay

Purified WT and K162C capsid particles in 1×PBS were divided into two aliquots and either left untreated or treated with 2 M sodium acetate pH 4.5. Samples were incubated on ice for 90 min, then plunge-frozen on graphene-oxide-coated Quantifoil copper mesh grids using a Vitrobot Mark IV (Thermo Fisher Scientific) at 4 °C and 100% relative humidity. Cryo-EM specimens were screened on a 200 kV FEG Glacios microscope (Thermo Fisher Scientific) equipped with a Falcon III direct electron detector.

### Quantifications and statistical analysis

No statistical methods were used to predetermine sample size, experiments were not randomized, and the investigators were not blinded to experimental outcomes. Data are represented as the mean of three independent experiments (**Fig. 4d**). Scatter plots were plotted with Microsoft Excel v.16. No statistical significance tests were used.

### Data availability

The atomic coordinates were deposited in the Protein Data Bank (www.rcsb.org) with accession codes XXXX [http://doi.org/10.2210/pdbxxxx/pdb], XXXX [http://doi.org/10.2210/pdbxxxx/pdb, XXXX [http://doi.org/10.2210/pdbxxxx/pdb, and XXXX [http://doi.org/10.2210/pdbxxxx/pdb]. The cryo-EM densities were deposited in the EM Data Bank with codes EMD-XXXXX, EMD-XXXXX, EMD-XXXXX, and EMD-XXXXX. Other source data are provided with this paper or from the corresponding author upon request.

## Acknowledgements

We thank the following facility staff for assistance in cryo-EM data collection: Bilal Ahsan, Giuseppe Cannone, Shaoxia Chen, Grigory Sharov, and other staff at the MRC-LMB EM Facility. We thank MRC-LMB Scientific Computing for computing support. We acknowledge the support of the MRC-LMB Media & Glass Wash facility. This work was supported by the Wellcome Trust [217191/Z/19/Z to Y.M.].

## Author Contributions

Conceptualization: HZ, YL, OBN, YM; Formal Analysis: HZ, YL, OBN, YM; Methodology: HZ, YL, OBN, YM; Investigation: HZ, YL, OBN, YM; Visualization: HZ, YM; Funding acquisition: YM; Project administration: YM; Supervision: YM; Writing – original draft: HZ, YM; Writing – review & editing: HZ, YM.

