## Supplementary Figures and Tables for "Capsid structures of Atlas viruses reveal an engineerable nanoparticle scaffold"

**Supplementary Table 1. Identification of the Atlas virus clade.**

| Name | GenBank accession | Species | Seq. cover[a] (%) | Seq. ID[b] (%) | Total score[b] | RMSD-Cα AF3 CA model vs. Atlas[c] (Å) | Seq. cover AF3 CA model vs. Atlas CA[c] (%) | Average pLDDT of AF3 model[d] |
|---|---|---|---|---|---|---|---|---|
| Atlas | EPB78661 | *A. ceylanicum* | 100 | 100 | 383 | N/A | 87 | 87 |
| Oos1 | KAK6016282 | *O. ostertagi* | 92 | 83 | 330 | 0.92 | 90 | 88 |
| Oos2 | KAK6031060 | *O. ostertagi* | 80 | 78 | 315 | 0.95 | 88 | 86 |
| Oos3 | KAK6026028 | *O. ostertagi* | 92 | 76 | 311 | 0.97 | 89 | 87 |
| Oos4 | KAK6028305 | *O. ostertagi* | 98 | 77 | 309 | 1.00 | 85 | 86 |
| Aca | RCN35992 | *A. caninum* | 93 | 67 | 272 | 0.86 | 87 | 88 |
| Ace2 | EPB75047 | *A. ceylanicum* | 99 | 67 | 264 | 0.96 | 87 | 87 |
| Nbr | VDL73942 | *N. brasiliensis* | 79 | 65 | 261 | 0.99 | 82 | 88 |
| Oos5 | KAK6030023 | *O. ostertagi* | 75 | 64 | 255 | 0.89 | 86 | 88 |
| Oos6 | KAK6018921 | *O. ostertagi* | 100 | 52 | 214 | 0.92 | 85 | 85 |
| Ace3 | EPB74949 | *A. ceylanicum* | 96 | 52 | 207 | 1.10 | 86 | 84 |

[a]PSI-BLAST against ClusteredNR database with Atlas virus polyprotein as query.

[b]BLASTp against the hits from the PSI-BLAST search with Atlas CA core structured region (a.a. 188–372) as query.

[c]An AlphaFold 3 (AF3) model of CA was superimposed on the Atlas CA AF3 model in UCSF Chimera[55]. The root mean square deviation (RMSD) and sequence cover were calculated after pruning with MatchMaker.

[d]The average predicted Local Distance Difference Test confidence score (pLDDT) was calculated prior to pruning with MatchMaker.

**Supplementary Table 2. Cryo-EM data collection, structure determination, and model refinement parameters.**

| | Atlas virus | | Atlas-Oos1 | Atlas-Nbr |
|---|---|---|---|---|
| **Data Collection and Processing** | | | | |
| Microscope/Detector | Krios/K3 | | Krios/K3 | Krios/K3 |
| Micrograph fluence ($e^{-}$ Å$^{-2}$) | 40 | | 40 | 40 |
| Exposure per frame ($e^{-}$ Å$^{-2}$) | 1.0 | | 1.0 | 1.0 |
| Nominal defocus range (µm) | -1.2 – -2.2 | | -1.2 – -2.2 | -1.2 – -2.2 |
| Pixel size (Å) | 0.826 | | 0.826 | 0.826 |
| N. movies in dataset | 23,292 | | 19,149 | 7,308 |
| N. initial particles picked | 150,458 | 9,969 | 356,523 | 209,204 |
| **Map averaging and refinement** | ***T* = 7** | ***T* = 12** | | |
| N. movies used | 15,279 | 4,065 | 6,451 | 7,007 |
| N. particles used | 32,697 | 5,408 | 8,891 | 151,130 |
| Symmetry | *I* (*T* = 7) | *I* (*T* = 12) | *I* (*T* = 4) | *I* (*T* = 1) |
| Map sharpening B factor (Å$^2$) | -98 | -98 | -76 | -81 |
| Resolution, unmasked ½-maps (FSC = 0.143) (Å) | 3.9 | 6.9 | 4.2 | 2.6 |
| Final resolution with masking (Å) | 3.2 | 4.4 | 3.5 | 2.2 |
| **Atomic model fit in data** | | | | |
| Map-model resolution (FSC 0.5, unmasked) | 3.3 | 6.3 | 3.7 | 2.5 |
| Map-model resolution (FSC 0.5, masked) | 3.3 | 5.9 | 3.7 | 2.4 |
| CC (mask), Phenix v2.0 | 0.82 | 0.74 | 0.84 | 0.91 |
| CC (volume), Phenix v2.0 | 0.82 | 0.75 | 0.83 | 0.91 |
| **Atomic model composition** | | | | |
| N. chains per ASU (*T* number) | 7 | 12 | 4 | 1 |
| N. non-hydrogen atoms | 10,115 | 17,365 | 5,844 | 1,453 |
| Protein residues | 1,288 | 2,211 | 744 | 181 |
| **Atomic model geometry, ADPs** | | | | |
| Bond lengths (Å) | 0.003 | 0.003 | 0.004 | 0.003 |
| Bond angles (˚) | 0.582 | 0.578 | 0.719 | 0.624 |
| Protein min/max/mean ADP | 44/165/89 | 103/170/125 | 66/117/84 | 42/145/75 |
| **Validation** | | | | |
| MolProbity score, Phenix v2.0 | 2.14 | 1.85 | 1.74 | 1.51 |
| ClashScore, Phenix v2.0 | 7.16 | 7.57 | 6.15 | 3.75 |
| Rotamer outliers (%) | 2.57 | 0.10 | 1.78 | 1.24 |
| Ramachandran plot | | | | |
| % favored | 93.3 | 93.3 | 96.7 | 96.1 |
| % allowed | 6.7 | 6.6 | 3.2 | 3.9 |
| % outliers | 0.0 | 0.1 | 0.1 | 0.0 |
| PDB code | 11XX | 11XX | 11XX | 11XX |
| EMDB code | EMD-00000 | EMD-00000 | EMD-00000 | EMD-00000 |

ASU, asymmetric unit.

ADPs, atomic displacement parameters.

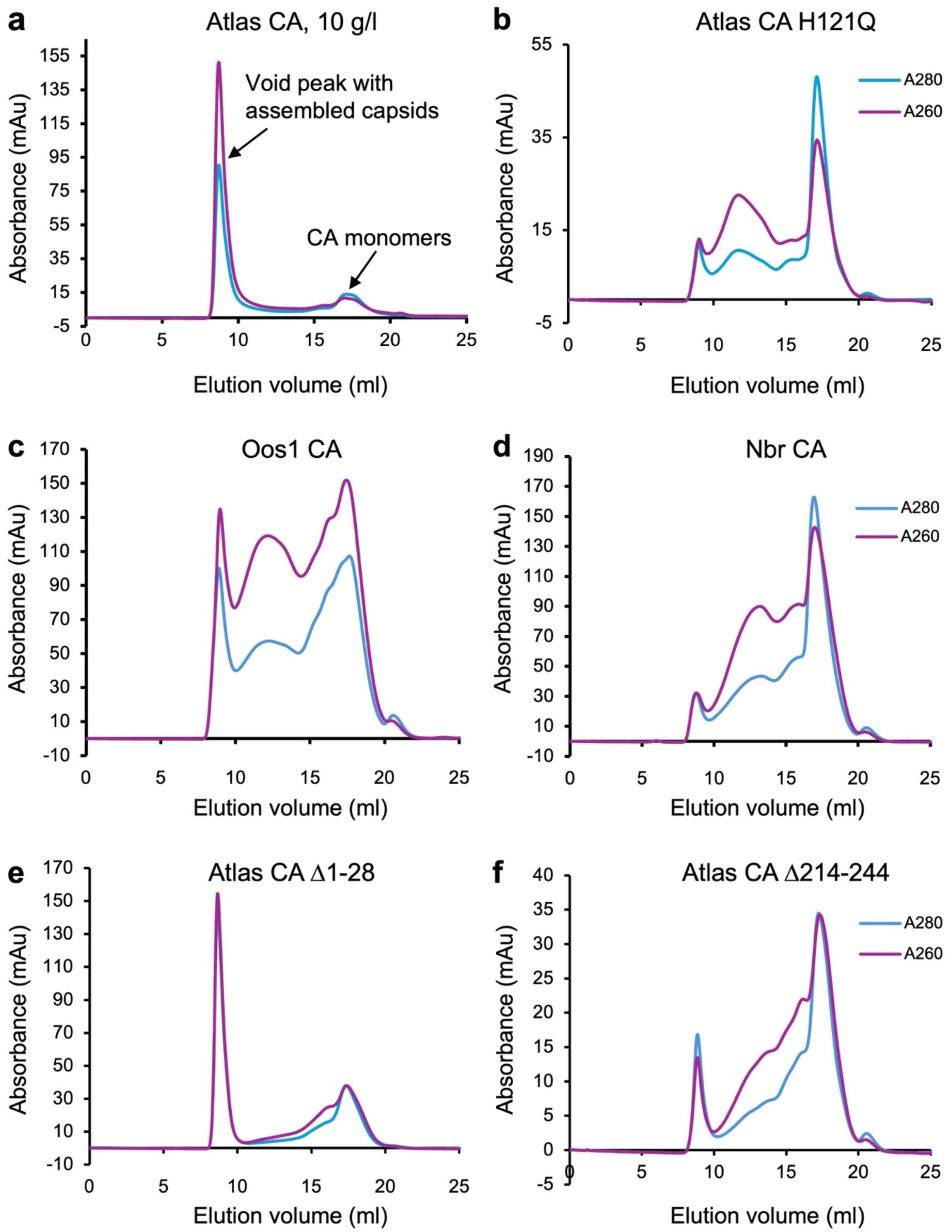


**Supplementary Fig. 1. Self-assembly of Atlas CA clade members and variants.** Size-exclusion chromatography profiles of selected members of Atlas-clade CA and indicated variants: (**a**) WT Atlas virus CA (same panel as Fig. 4a); (**b**) H121Q Atlas virus CA; (**c**) Atlas-Oos1 CA; (**d**) Atlas-Nbr CA; (**e**) Atlas virus CA N-terminal tail deletion (Δ1-28); (**f**) Atlas virus CA C-terminal tail deletion (Δ214-244). All samples were analyzed at 7-10 g/l protein concentration in 2×PBS, 1 M NaCl, and 1 mM DTT.

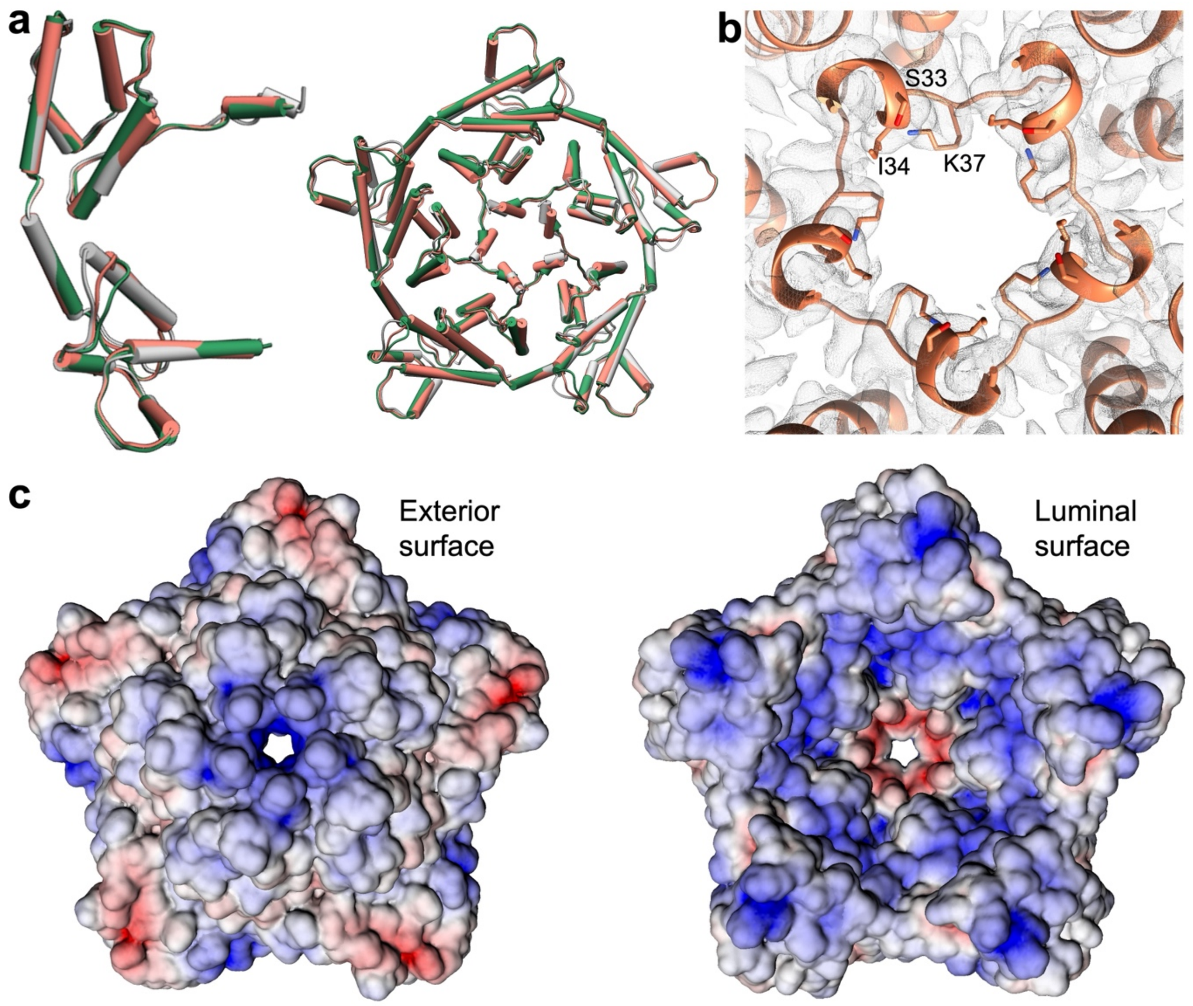


**Supplementary Fig. 2.** Structures and properties of pentameric capsomeres. (**a**) Superposition of capsid protomers (left) and pentameric capsomeres (right) from Atlas virus *T* = 7 (pink), Atlas-Oos1 (green), and Atlas-Nbr (grey) particles. (**b**) Density map and atomic model of the Atlas-Nbr pentamer core. Side chains of the pore-lining residues (Asp31, Ser33, Ile34, and Lys37; corresponding to Asp35, Ser37, Leu38, and Lys41 in Atlas CA) are shown in stick representation. (**c**) Solvent-accessible surface representations of the Nbr pentamer viewed from the capsid exterior (left) and lumen (right). Surfaces are colored by electrostatic potential (APBS, https://server.poissonboltzmann.org)[58], ranging from -5 (red, acidic) to +5 (blue, basic) kT/e.

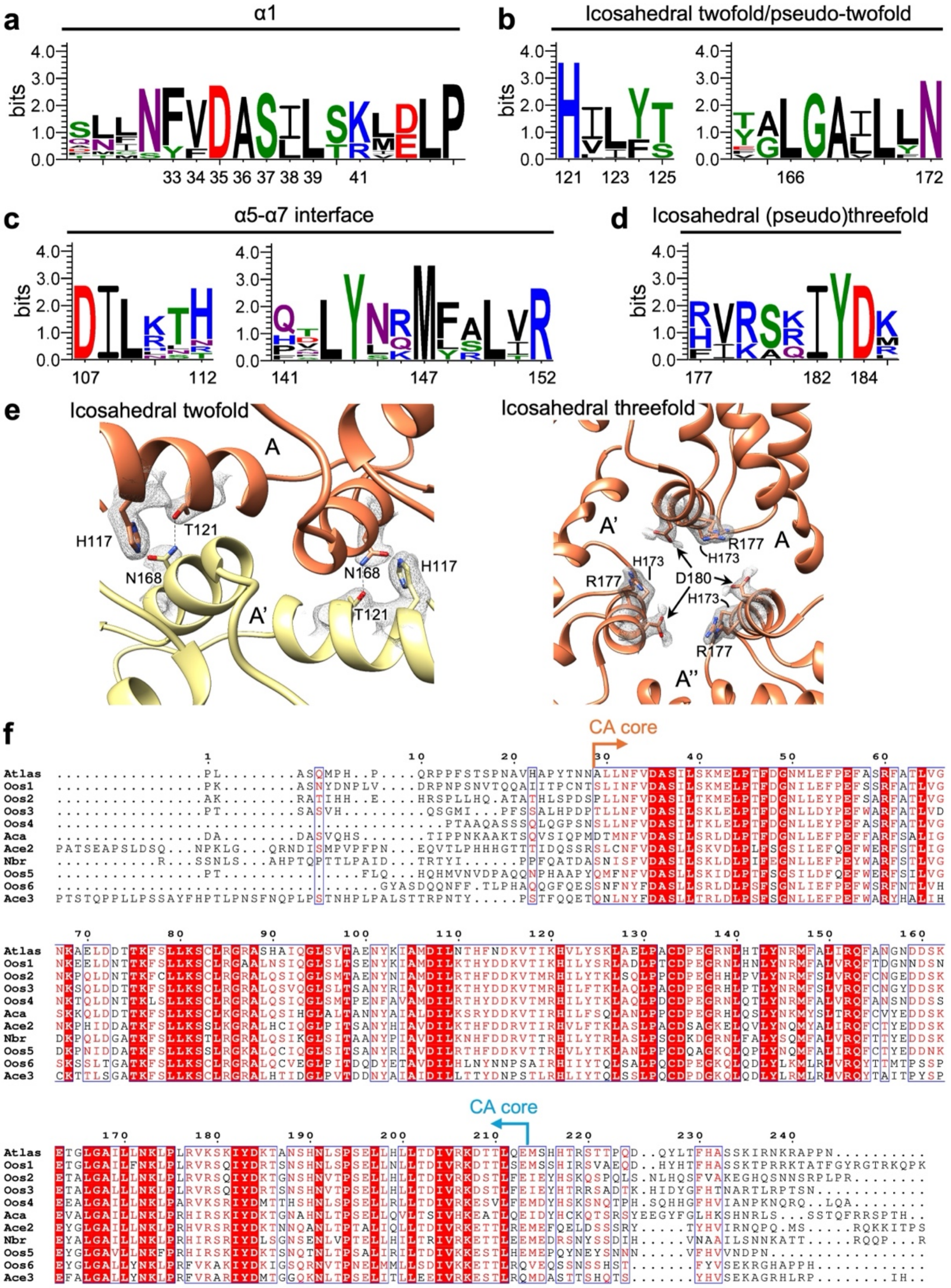


**Supplementary Fig. 3. Capsid sequence conservation in the Atlas clade.** (**a-d**) Sequence logos were generated with WebLogo3[53] from a multiple sequence alignment (MSA) of 11 CA orthologs. Residue numbers refer to the Atlas virus CA sequence. The y-axis represents information content in bits. (**a**) Helix α1. Key structural positions are indicated: the hydrophobic core (residues 33, 34, 36, 38, 39), an exterior-facing electronegative residue (Asp35), a pore-lining polar ring (Ser37), and a lumen-facing electropositive ring (Lys/Arg41). (**b**) Twofold and pseudo-twofold axes. Intercapsomere hydrogen-bonding residues identified in Atlas virus CA are shown. (**c**) Intracapsomere α5–α7 interface, highlighting the 107–152 salt bridge and the less conserved 112–141 hydrogen-bonding pair. (**d**) Threefold and pseudo-threefold axes, featuring a partially conserved basic patch (177, 181, 185) and acidic residue (Asp184). (**e**) Closeups of the twofold- and threefold-axis interfaces of the *T* = 1 Atlas-Nbr structure. (**f**) Complete MSA of the 11 CA sequences (Clustal Omega[51], rendered with ESPript 3.2[52]). Arrows indicate the boundaries of the structured CA core region.

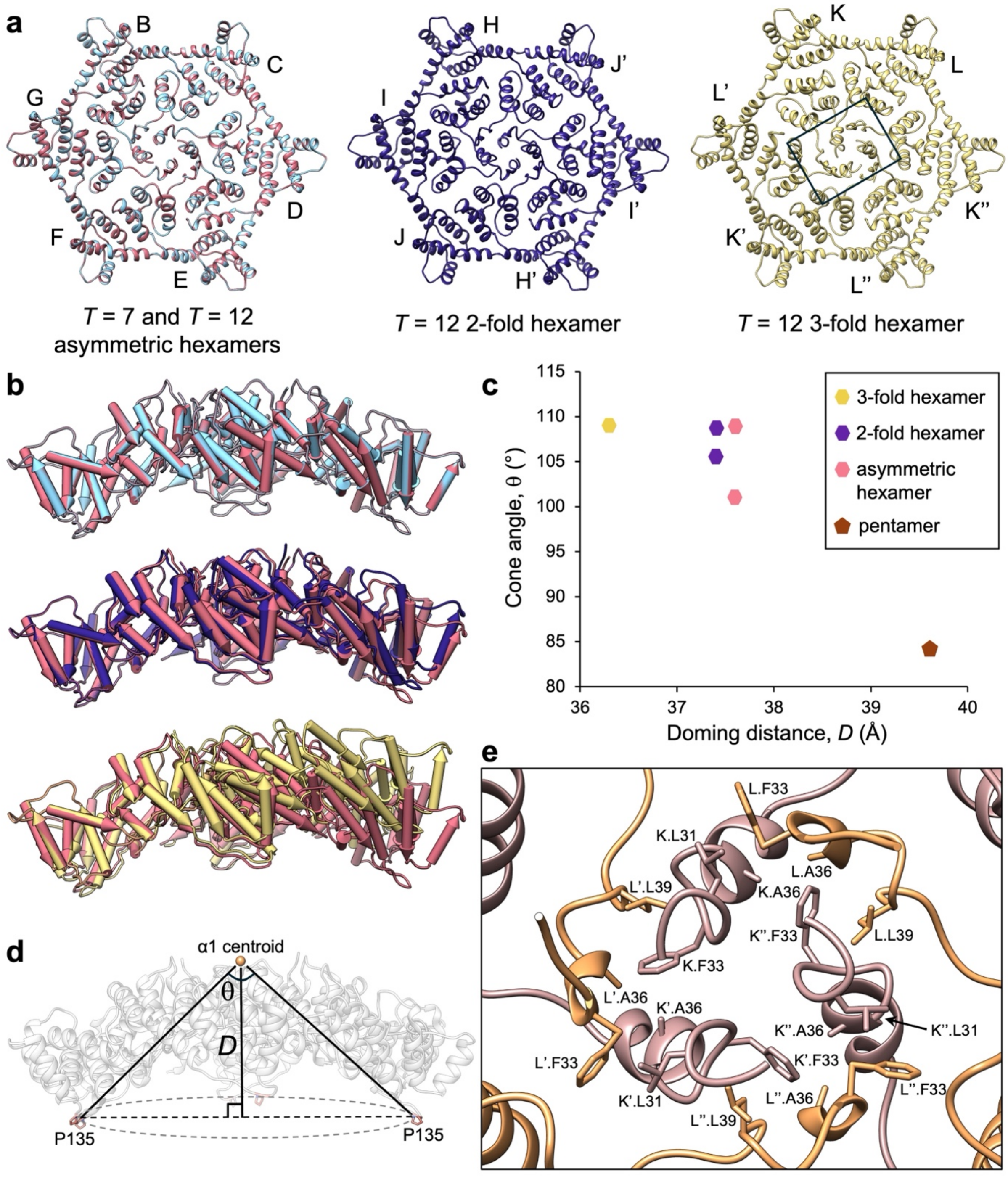


**Supplementary Fig. 4. Structural plasticity of capsid hexamers.** (**a**) Models of the different Atlas virus hexamers: *T* = 7 asymmetric (light blue), *T* = 12 asymmetric (pink), *T* = 12 twofold-symmetric (purple), and *T* = 12 threefold-symmetric (gold). Chain identifiers correspond to Fig. 2, with symmetry-related copies denoted by prime (') and double prime ("). (**b**) Side views of superimposed hexamers: *T* = 7 and *T* = 12 asymmetric (top); *T* = 12 asymmetric and twofold-symmetric (middle); *T* = 12 asymmetric and threefold-symmetric (bottom). (**c**) Quantification of hexamer curvature in terms of hexamer cone angle versus doming distance, as defined in *(d)*. (**d**) The cone angle (θ) is the angle formed by lines from a Pro135 Cα atom to the α1 centroid to the Pro135 Cα atom of the opposing chain. The doming distance, *D*, is the distance from the α1 centroid to the plane defined by the P135 Cα atoms in the capsomere. (**e**) Structure of the threefold-symmetric hexamer core (closeup of area boxed in *(a)*).

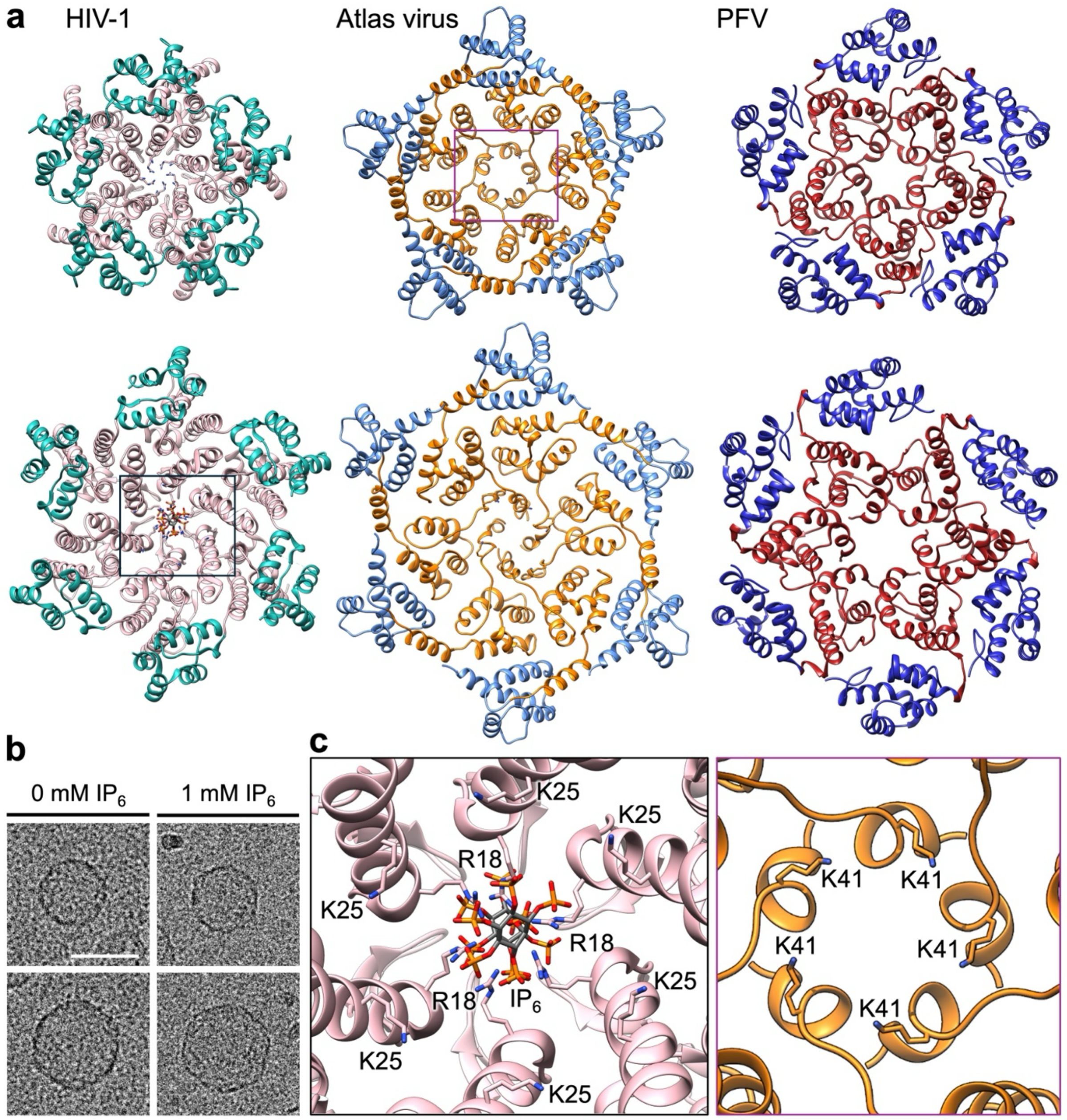


**Supplementary Fig. 5. Structural comparison of HIV-1, Atlas, and prototype foamy virus (PFV) capsomeres.** (**a**) Luminal views of the HIV-1 mature capsid pentamer (PDB 8CKW) and inositol hexaphosphate ($IP_6$)-bound hexamer (PDB 6BHT)[42]; the Atlas virus $T$ = 7 pentamer and hexamer; and the PFV pentamer (PDB 8OZL)[33] and hexamer (PDB 8OZM)[33]. (**b**) Cryo-EM micrographs of $T$ = 7 (top) and $T$ = 12 (bottom) Atlas virus particles assembled without (left) or with (right) 1 mM $IP_6$. Scale bar, 50 nm. (**c**) Closeup views of the capsomere pores (closeups of the areas boxed in *(a)*). Left, the $IP_6$ coordination site in the HIV-1 mature hexamer, where two $IP_6$ molecules are coordinated by stacked rings of Arg18 and Lys25 side chains. Right, the Atlas virus pentamer pore, highlighting the pore-lining Lys41 side chains.

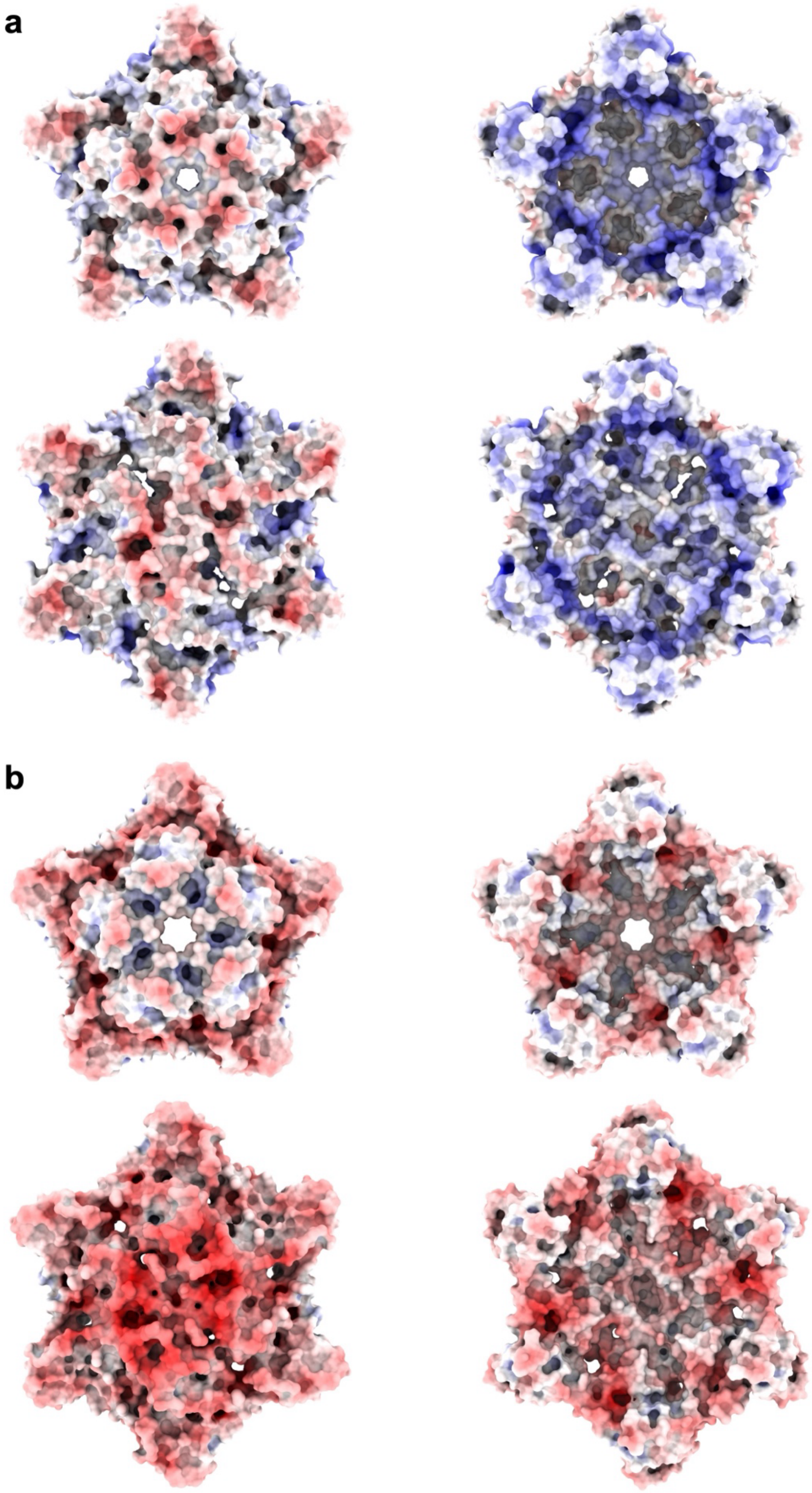


**Supplementary Fig. 6. Electrostatic surface potentials of Atlas-clade capsomeres.** Solvent excluded surfaces of capsomeres from Atlas virus, (**a**), and Atlas-Oos1, (**b**). Left, capsomeres viewed from the particle exterior side. Right, view from the particle lumen. Surfaces are colored by electrostatic potential calculated with APBS[58], ranging from -10 (red, acidic) to +10 (blue, basic) kT/e. Images drawn with ChimeraX[59].
